# D2 autoreceptors gate vulnerability to cocaine use disorder

**DOI:** 10.64898/2026.03.10.710882

**Authors:** Erin M Murray, Daniel Diaz-Urbina, Roland Bock, Emilya Ventriglia, Anna Tischer, Jung Hoon Shin, Seul Ah Lee, Lucy G Anderson, Sydney Cerveny, Isabel Bleimeister, Miriam E Bocarsly, Michael Michaelides, Veronica A Alvarez

## Abstract

A defining feature of substance use disorder is that repeated drug use does not always lead to addiction, motivating the search for biomarkers of vulnerability^1^. Reduced striatal dopamine D2/3 receptor availability is a robust PET correlate of problematic stimulant use^2–5^, but the signal may reflect high endogenous dopamine level, and it conflates presynaptic D2 autoreceptors on dopamine axons with postsynaptic D2/3 heteroreceptors on striatal projection neurons. We dissociated these contributions using cell type–specific Drd2 haploinsufficiency in dopamine neurons (autoD2KD), D2-expressing medium spiny neurons (MSN-D2KD), or both. Autoreceptor haploinsufficiency (autoD2KD) weakened presynaptic control of dopamine release, enhanced phasic gain, and prolonged cocaine-evoked dopamine elevations. This was accompanied by a hyper-exploratory trait and altered cocaine adaptation. Specifically, autoD2KD mice showed greater cocaine-seeking behavior, despite intact responses to sucrose reward and punishment. Although all genotypes showed graded reductions in striatal D2/3 binding, D1-like compensations diverged, resulting in different D1:D2/3 ratio in the striatum. The clinical implication is that striatal D1 density and D1:D2/3 balance may emerge as critical biomarkers for distinguishing cell-type-specific D2 reductions relevant to addiction vulnerability.

## INTRODUCTION

Substance use disorder (SUD) is a chronic, relapsing condition characterized by compulsive drug seeking and use despite adverse consequences. The DSM-5-TR defines SUD using 11 criteria spanning four domains—impaired control, social impairment, risky use, and tolerance/withdrawal—many of which can be operationalized in rodents^6–9^. Yet diagnostic criteria do not capture a defining epidemiological feature of addiction: repeated exposure does not inevitably lead to disorder^1^. With cocaine, for example, most exposed individuals do not develop cocaine use disorder^10^. This variability motivates two translational questions: which biological factors confer vulnerability to compulsive drug seeking and taking, and which biomarkers could identify high-risk individuals before persistent use emerges?

Vulnerability to stimulant use disorder is shaped by psychiatric comorbidities and individual differences in behavioral traits, which arise from interacting biological and environmental influences. One candidate mechanism linking SUD vulnerability and psychiatric comorbidity to sustained drug use is the self-medication hypothesis, which proposes that some individuals continue using because the drug transiently normalizes an aversive internal state or improves functioning, thereby reinforcing repeated use despite long-term costs^7,8^. In this framework, trait-like differences in novelty seeking, exploration, and risk processing could bias trajectories toward continued use by increasing either the reinforcing value of drug effects or the perceived functional benefits of intoxication^11,12^.

Reduced striatal dopamine D2/3 receptor availability is among the most replicated PET findings in stimulant use disorders^2–5,13^. Mechanistic interpretation is limited for several reasons. First, commonly used ligands bind to D2/3-family and may reflect contributions from both D2 and D3 receptors and endogenous levels^14,15^. Second, lower D2/3 availability may reflect pre-existing vulnerability, drug-induced adaptations, or both^4,16–19^. Third—and central to the present study—the cellular sources of the PET signal, and thus the relevant circuit mechanisms and therapeutic targets, remain unresolved.

The striatum expresses high levels of both D1- and D2-family dopamine receptors. D1 receptors are enriched in direct-pathway projection neurons, whereas striatal D2/3 PET ligand binding reflects receptors in at least two functional compartments: postsynaptic D2-family receptors (heteroreceptors) on striatal GABAergic projection neurons and presynaptic D2 autoreceptors on dopaminergic axons arising from midbrain neurons that innervate the striatum^20^. Postsynaptic D2 signaling modulates basal ganglia output and reinforcement-related learning, whereas D2 autoreceptors provide feedback inhibition that constrains dopamine synthesis, release and neuronal firing^21–25^. Reduced striatal D2/3 PET signal could therefore reflect hypofunction in one or both compartments, with distinct circuit consequences and implications for stimulant vulnerability.

We hypothesized that reduced postsynaptic D2 heteroreceptor signaling increases the effective D1/D2 balance in striatal output pathways, enhancing susceptibility to stimulant-induced plasticity and reinforcement^26–28^. Alternatively, reduced D2 autoreceptor function is hypothesized to amplify stimulant-evoked dopamine signaling by weakening presynaptic feedback inhibition, thereby increasing cocaine sensitivity and progression to compulsive use^7,18,24,25,29^. A third hypothesis is that concurrent reductions in autoreceptors and heteroreceptors interact to reshape dopamine dynamics and striatal output in ways not predictable from either perturbation alone.

## RESULTS

To dissociate and test these hypotheses in vivo, we generated mice with cell-type–specific Drd2 haploinsufficiency: knockdown of D2 autoreceptors in midbrain dopamine neurons (autoD2KD), knockdown of D2 heteroreceptors in D2-expressing medium spiny neurons (MSN-D2KD), or both (double-D2KD), with littermate controls. We quantified dopamine-dependent baseline traits (novelty-driven exploration and approach–avoidance behavior) and assessed cocaine-relevant phenotypes spanning acute and repeated drug responses and DSM-5-TR–aligned SUD-like behaviors that can be operationalized in rodents, including high-effort consumption, perseverative seeking during drug unavailability, punishment-resistant intake, and abstinence-related seeking/craving^9,12,30–32^. The results would inform us whether D2 autoreceptors or heteroreceptors are determinant of cocaine vulnerability and suggest that striatal D1 binding—and the D1:D2/3 balance—may help distinguish the compartment-specific biological substrates underlying low striatal D2/3 PET signal.

### Generation of cell-type-specific D2 receptor knockdown mice

To achieve cell-type-specific reductions in D2 receptors, Drd2 gene expression was selectively downregulated in dopamine neurons, striatal medium spiny neurons (MSNs), or both populations simultaneously, generating male and female mice with reduced autoreceptors (autoD2KD), heteroreceptors (MSN-D2KD), or both (double-D2KD) (Fig. 1a,b). The breeding strategy yielded littermates of all four genotypes in approximately Mendelian proportions (25.2% WT, 26.8% autoD2KD, 23.5% MSN-D2KD, 25.2% double-D2KD; n = 941 mice; Chi-square test χ²(3) = 1.64, p = 0.65), indicating no gross effects on viability or development. Validation of Drd2 knockdown was performed using quantitative PCR to measure Drd2 mRNA levels in striatum, with cortex serving as a control region (Fig. 1b). Drd2 mRNA was reduced in both nucleus accumbens (NAc) and dorsal striatum for MSN-D2KD and double-D2KD mice, whereas autoD2KD mice showed no change (Fig. 1b). Analysis revealed main effect of region and genotype (region: F (1.25, 17.46) = 5.34, p = 0.027; genotype: F (3, 16) = 12.81, p = 0.0002) with post hoc comparisons confirming reductions in MSN-D2KD (NAc: p = 0.019, dStr: p = 0.0002) and double-D2KD (NAc: p = 0.021; dStr: p = 0.001) and no change in autoD2KD (NAc: p = 0.48; dStr: p = 0.302) when compared to controls.

**Figure 1.**
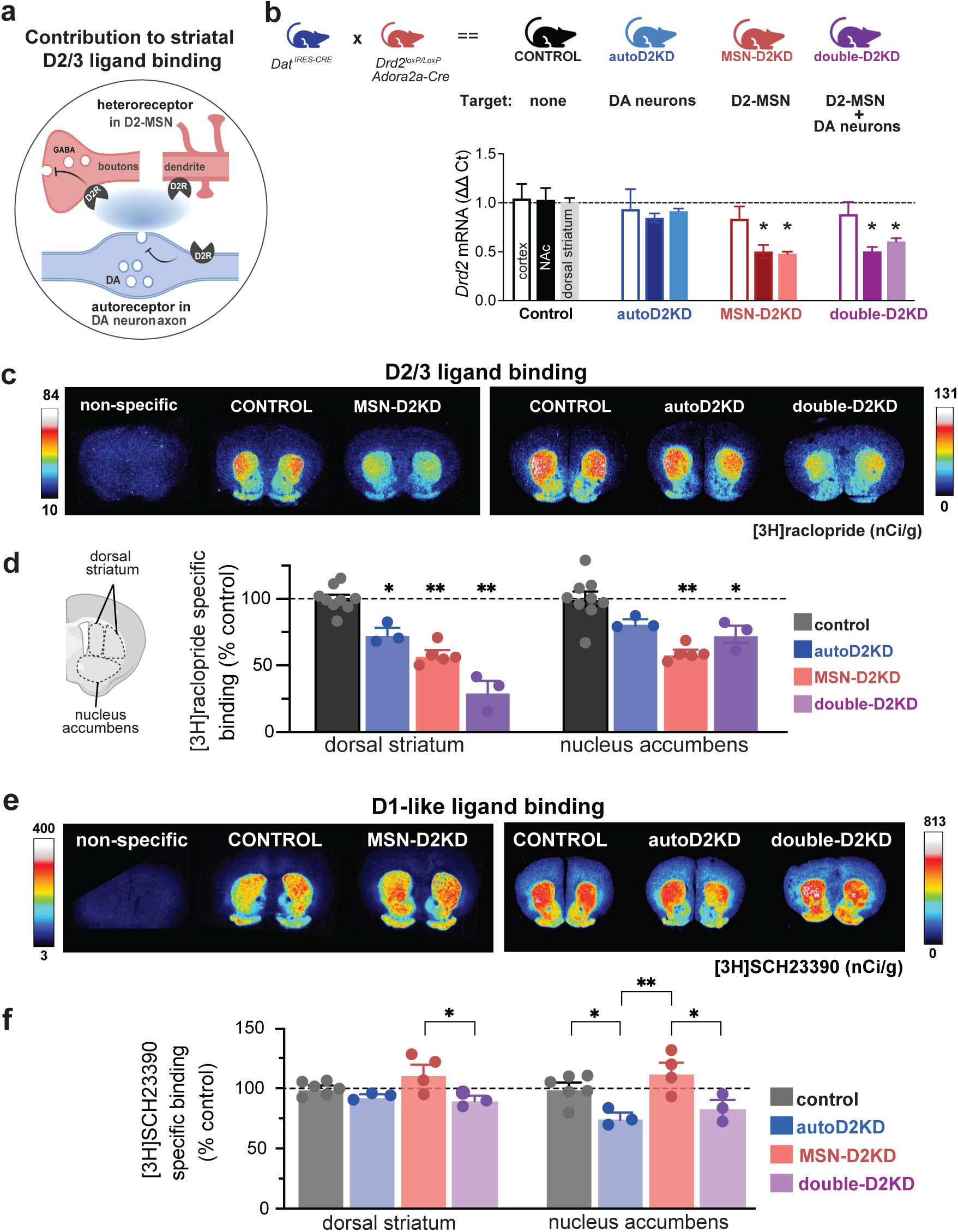
Cell-type–specific Drd2 haploinsufficiency produces graded D2/3 reductions and divergent D1 remodeling. **a**, Schematic of the principal contributors to striatal D2/3 ligand binding: postsynaptic D2 heteroreceptors on medium spiny neurons (MSNs; red) and presynaptic D2 autoreceptors on dopamine axons (blue). **b**, Breeding strategy generating littermates with cell-type–specific Drd2 haploinsufficiency. Genotypes: control (Drd2^loxP/wt^; black), autoD2KD (Drd2^loxP/wt^; Dat^IRES-Cre/wt^; blue), MSN-D2KD (Drd2^loxP/wt^; Adora2a-Cre^+/–^; red), and double-D2KD (Drd2^flox/wt^; Adora2a-Cre^+/–^; Dat^IRES-Cre/wt^; purple). Bottom, RT–qPCR quantification of Drd2 mRNA (ΔΔCt) in frontal cortex (open), nucleus accumbens (NAc; dark), and dorsal striatum (light) (n = 5 control; n = 3–5 autoD2KD; n = 5 MSN-D2KD; n = 3–5 double-D2KD). **c, e**, Representative autoradiographs of [3H]raclopride binding (c) and [3H]SCH23390 binding (e), including nonspecific binding. Color scale indicates nCi/g. Autoradiography was performed in two independent cohorts, each with its own control group; representative control images are therefore shown twice. **d, f**, Left, schematic of striatal regions of interest used for quantification. Right, specific binding in dorsal striatum and NAc, normalized to the mean of the corresponding control region, for [3H]raclopride (d) and [3H]SCH23390 (f). Symbols denote individual mice; bars show mean ± s.e.m. For binding quantification, n = 3–9 mice per genotype; 8–12 sections per mouse; 24–30 ROIs per animal. *P ≤ 0.05; #0.1 ≥ P > 0.05.

### Cell-type–specific Drd2 haploinsufficiency yields graded reductions in striatal D2/3 binding

To quantify the impact of cell-type specific Drd2 knockdown on D2 receptor protein levels, we measured D2/3 ligand binding in brain sections from all four genotypes using [^3^H]raclopride—the same ligand class used in human PET studies. Autoradiography revealed significant reductions in D2/3 binding across both the dorsal striatum and NAc in all three knockdown genotypes relative to littermate controls (Fig. 1c-d). However, the magnitudes differed by genotype and region. In the dorsal striatum, double-D2KD mice exhibited the largest reduction in D2/3 binding (70 ± 8%), followed by MSN-D2KD (42 ± 3%) and autoD2KD mice (26 ± 4.5%). In the NAc, MSN-D2KD and autoD2KD mice showed reductions comparable to those observed in the dorsal striatum (41 ± 2.6% and 18 ± 2.6%, respectively). In contrast, double-D2KD mice showed a markedly smaller decrease in the NAc (27 ± 6%) than the dorsal striatum. Accordingly, a significant Genotype × Region interaction was detected (F(3,32) = 5.86, p = 0.0026). Thus, combined targeting of D2 autoreceptors and heteroreceptors produced the greatest reduction in dorsal striatal D2/3 binding, whereas the largest reduction in NAc occurred when heteroreceptors alone were targeted. These graded, region- and cell-type-specific reductions in striatal D2/3 binding enable us to test how these distinct dopaminergic perturbations translate into behavioral traits and vulnerability to cocaine-related SUD phenotypes.

### Genotype-specific remodeling of striatal D1-like ligand binding that reshapes D1:D2/3 balance

Autoradiography with [^3^H]SCH-23390 revealed region- and genotype-dependent differences in D1-like binding (genotype x region F(3,12) = 5.1 p = 0.016; genotype F(3,12) = 5.9, p = 0.009; region F(1,12) = 10.7 p = 0.006). D1-like ligand binding was reduced in the NAc of autoD2KD mice compared to controls (p = 0.01; Fig. 1e-f). Double-D2KD mice showed no significant change from controls (p = 0.16) or autoD2KD (p = 0.797). By contrast, MSN-D2KD mice had enhanced NAc D1-like binding than autoD2KD and double-D2KD mice (p = 0.0006 and p = 0.007, respectively); while remaining comparable to controls (p = 0.24; Fig. 1f).

Dopamine-dependent behaviors are strongly shaped by the relative density and activity of dopamine D1- versus D2-like receptors, suggesting that D1:D2/3 binding ratio may represent a compact readout of the balance between these receptor classes across regions. To this end, we computed a D1:D2/3 binding ratio for each striatal subregion, defined as the ratio of [^3^H]SCH-23390 binding to [^3^H]raclopride binding (Fig. 1g). This revealed a genotype effect and a genotype x region interaction (F(3,25) = 22.7, p = 0.0001; F(3,25) = 7.26, p = 0.001). In autoD2KD mice, the D1:D2/3 binding ratio was unchanged compared to controls in both the dorsal striatum (p = 0.483) and NAc (p = 0.758). Despite reduced D2/3 binding, a proportional decrease in D1 binding preserved the overall balance (Fig. 1h).

In contrast, MSN-D2KD mice showed a robust increase in the D1:D2/3 binding ratio in both the dorsal striatum (p < 0.0001) and nucleus accumbens (p = 0.0002), consistent with preserved (or modestly increased) D1 binding coupled to pronounced reductions in D2/3 binding^27^ (Fig. 1h). Double-D2KD mice exhibited a regionally selective shift, with elevated ratios in dorsal striatum (p < 0.0001, Fig. 1h) but not in nucleus accumbens (p = 0.884, Fig. 1h), likely driven by the comparatively smaller reduction in accumbens D2/3 binding (Fig. 1). Together, these data indicate that autoD2KD mice maintain striatal D1:D2/3 balance via coordinated downregulation of D1 binding, whereas MSN-D2KD mice show a broad, striatum-wide skew toward higher D1 relative to D2/3 binding.

### Preserved D2/3 agonist responses with genotype-specific alterations in D1-like signalling

In controls, systemic D2/3 agonist quinelorane (5–10 μg/kg, i.p.) suppressed locomotion in a dose-dependent manner (One-way ANOVA: F(1.954, 8.794) = 12.46, p = 0.003; Supplementary Fig. 1). As expected, this suppression was markedly attenuated in mice with complete loss of D2 autoreceptors (auto-D2KO; genotype: F(1,18) = 4.34, p = 0.052), validating assay sensitivity (Bello et al., 2011; Anzalone et al., 2012). In contrast, quinelorane-induced suppression was preserved in all three Drd2 knockdown genotypes and did not differ from controls (5 μg/kg: F(3,31) = 0.66, p = 0.583; Figure 2 c,d; Supplementary Fig. 1), indicating intact behavioral responsiveness to systemic D2/3 agonism despite reduced ligand binding.

**Figure 2.**
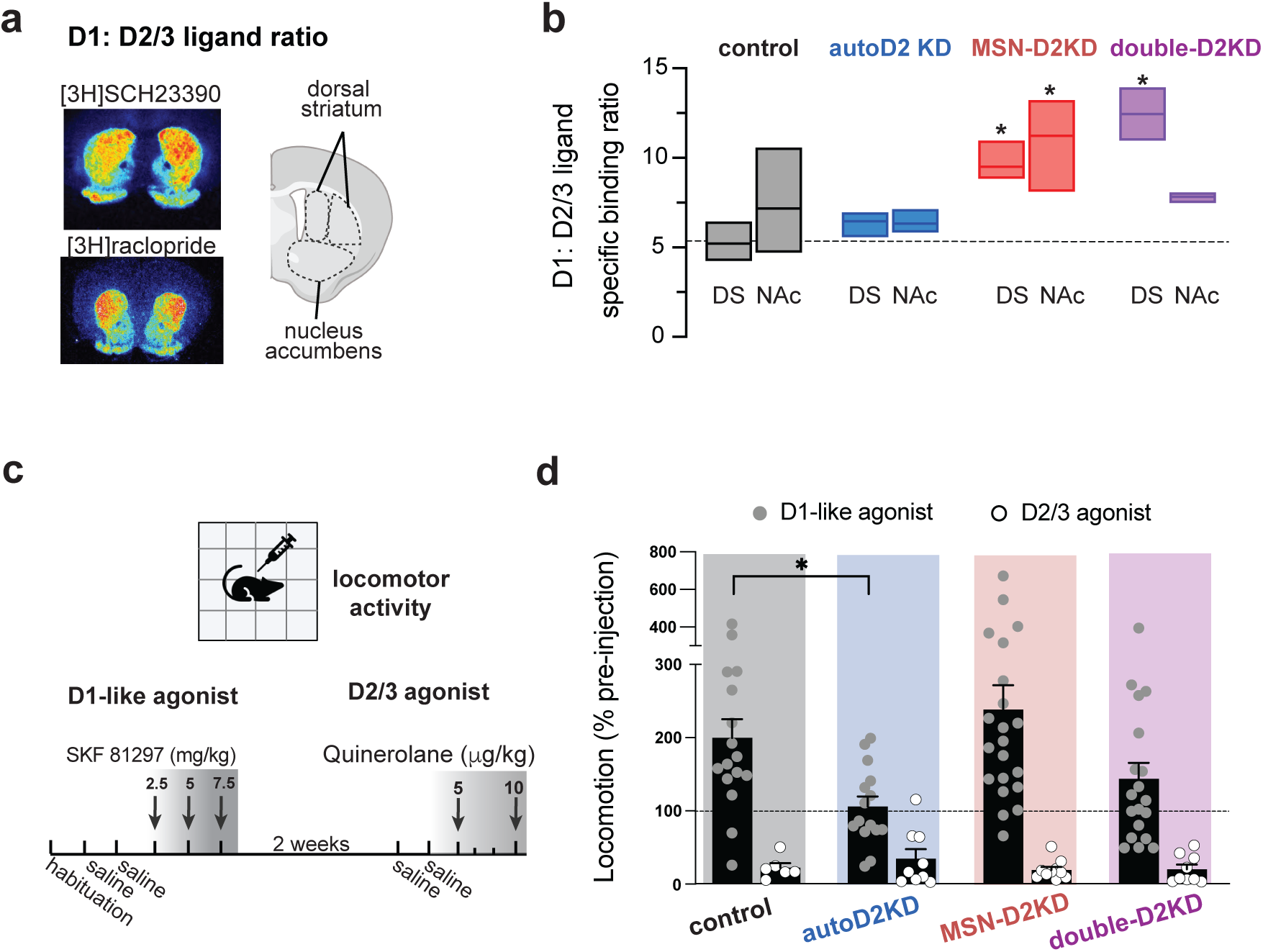
Striatal D1:D2/3 binding ratio and agonist-mediated locomotor responses. **a,** Representative autoradiographs showing D1-like ligand binding ([3H]SCH93330) and D2/3 ligand binding ([3H]raclopride) for a control mouse with cartoon indicating quantified striatal subregions (dorsal striatum and nucleus accumbens). **b,** D1:D2/3 ratio ([3H]SCH93330/[3H]raclopride) quantified for each region across genotypes (n=3-6 mice per region, per genotype). Box plots indicate mean and min-max range. Dash line marks mean ratio in the dorsal striatum for control mice. *p ≤ 0.05 versus control. **c,** Timeline for assessing locomotor responses to dopamine receptor agonists. After habituation and saline injections (2 d), mice received escalating doses of the D1-like agonist SKF81297 (2.5, 5, 7.5 mg/kg, i.p.). After a 2-week washout, mice received saline (2 d) followed by the D2/3 agonist quinelorane (5 and 10 μg/kg, i.p.). **d**, Locomotor responses normalized to each mouse’s pre-injection baseline, shown after SKF81297 (5 mg/kg; filled symbols) and quinelorane (5 μg/kg; open symbols). Points indicate individual mice; bars show mean ± s.e.m. For all panels, *p ≤ 0.05 versus control.

In contrast, locomotor responses to the D1-like agonist SKF81297 (2.5–7.5 mg/kg, i.p.) differed by genotype (genotype: F(4,80) = 6.36, p = 0.025; dose: F(2.373,178.8) = 46.98, p < 0.0001; genotype × dose: F(12,226) = 2.00, p = 0.025; Supplement Figure 1). At 2.5 mg/kg, SKF81297 approximately doubled locomotion in controls (199 ± 26%) but was attenuated in autoD2KD mice (106 ± 14%; Tukey p = 0.03) and potentiated in MSN-D2KD mice (238 ± 33%) relative to autoD2KD and double-D2KD mice (144 ± 22%; p = 0.0003 and 0.009) (Fig. 2b; Supplementary Fig. 1). Across agonists, we observed a significant agonist × genotype interaction (F(3,97) = 2.89, p = 0.039; agonist: F(1,97) = 56.21, p = 0.0001; Fig. 2d). Because systemic agonist responses can be buffered by circuit-level compensation—and D2/3 binding reductions were modest in autoD2KD mice—we next assessed presynaptic control of dopamine release and cocaine-evoked dopamine dynamics directly in these mice.

### Phasic dopamine is potentiated and cocaine effects are prolonged in mice with reduced autoreceptor function

In *ex vivo* striatal recordings, the D2/3 agonist quinpirole (30 nM) suppressed evoked dopamine signals less in autoD2KD mice than in controls (t(20) = 2.87, p = 0.009; Fig. 3a-b). Dopamine release evoked by low-frequency stimulation (single pulse delivered every 60s; 0.016Hz) was unchanged (controls 13.0 ± 1.5 μM; autoD2KD 13.7 ± 1.7 μM; t(22) = 0.21, p = 0.83), whereas high-frequency stimulation (20 Hz train) elicited a larger response in autoD2KD mice when expressed relative to tonic release (train/single = 4.5 ± 0.5 versus 3.8 ± 0.5; t(22) = 3.44, p = 0.002; Fig. 3c). Acute cocaine increased evoked dopamine signals and slowed their decay in both genotypes (Fig. 3d-e) but the time course differed by genotype (time x genotype interaction χ2 = 373, p < 0.0001). In controls, peak amplitude declining over time, whereas in autoD2KD mice potentiation persisted, yielding larger signals after prolonged exposure (at 40 min: 131 ± 13% versus 91 ± 6% of baseline in controls; t(16) = 2.7, p = 0.017; Fig. 3f). Together, these data indicate that reduced D2 autoreceptor function enhances phasic dopamine gain and prolongs cocaine-evoked dopamine elevations.

**Figure 3.**
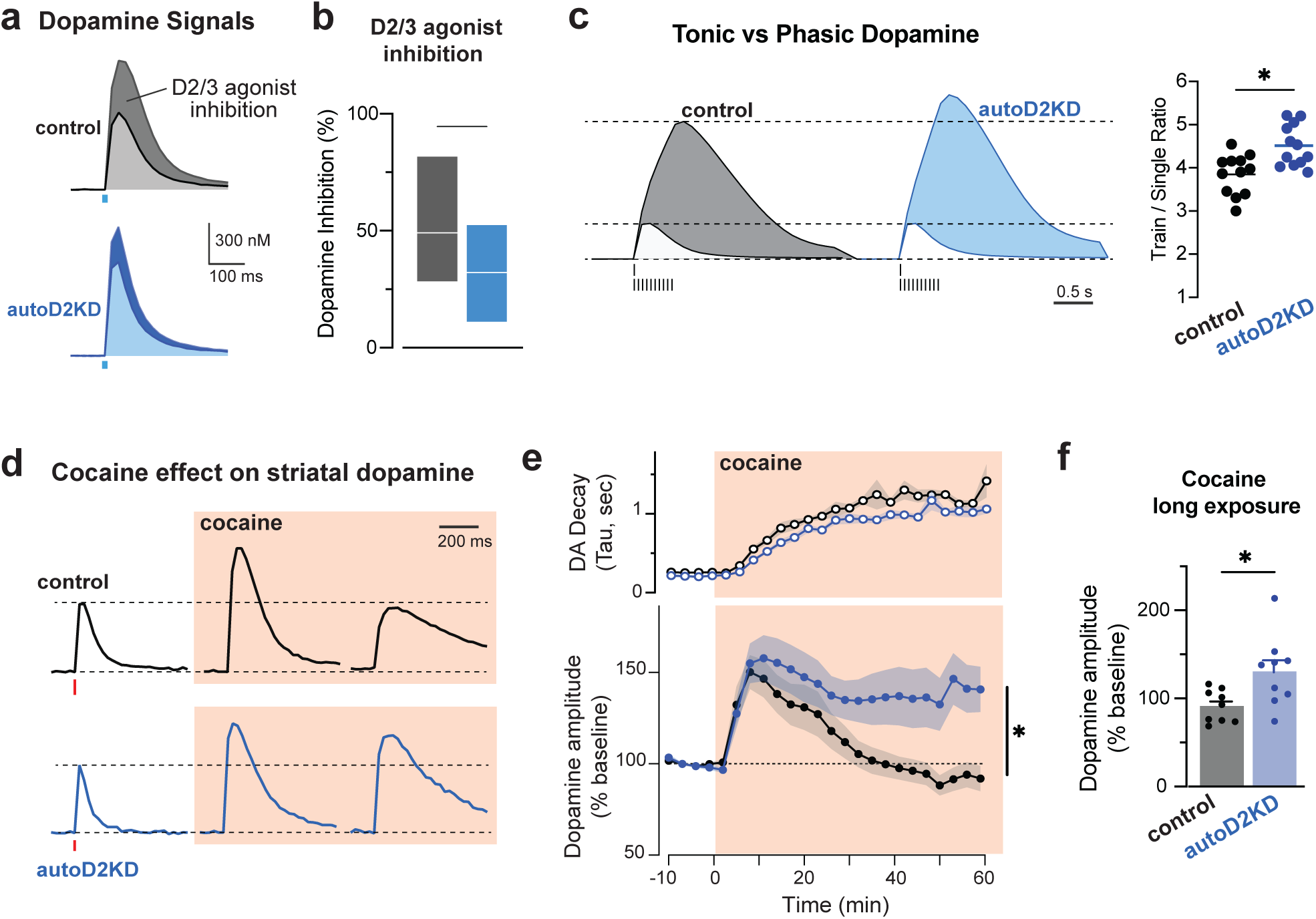
Reduced autoreceptor-mediated feedback increases phasic dopamine gain and prolongs cocaine-evoked dopamine elevations. **a**, Representative striatal fast-scan cyclic voltammetry traces evoked by electrical stimulation in control Dat^IRES-Cre^ mice (black) and autoD2KD mice (blue) before (dark) and after (light) bath application of the D2/3 agonist quinpirole (30 nM). **b**, Quinpirole-induced inhibition of evoked dopamine release, expressed as percent suppression from baseline (control: n = 12 slices from 4 mice; autoD2KD: n = 10 slices from 4 mice). *p = 0.009. Box plots show min-max range and mean in white line. **c**, Phasic-to-tonic gain. Left, representative dopamine traces evoked by a single pulse (light) or a 20 Hz train of 10 pulses (dark), scaled to the single-pulse peak. Right, amplitude ratio of train-evoked relative to single-pulse evoked (control and autoD2KD: n = 12 slices from 2 mice per genotype). Symbols are individual slice data and lines indicate means. *p = 0.002. **d**, Representative dopamine transients evoked by single-pulse stimulation at baseline and after cocaine (3 μM; orange shading) for 6 and 60 min. **e**, Time course of cocaine effects on dopamine signal decay (top; τ) and peak amplitude (bottom; normalized to pre-cocaine baseline) in control (black) and autoD2KD (blue) slices. *p<0.0001 Cocaine Timecourse x Genotype interaction. **f**, Dopamine peak amplitude after 60 min cocaine, normalized to baseline (n = 9 slices from 3 mice per genotype). *p = 0.017.

### Opposite shifts in exploratory traits among mice with D2 knockdowns

We predicted that altering phasic dopamine responses or the relative signaling via D1 and D2/3 receptors would modify dopamine-dependent traits, including novelty-driven exploration and risk avoidance. Novelty-induced exploration was quantified in an open field, which revealed a significant effect of genotype (one-way ANOVA: F(3, 183) = 17.28, p < 0.0001; Fig. 4a,c, Supplementary Fig 1). Compared to controls, the distribution of novelty-induced locomotion was shifted leftward in MSN-D2KD mice (p = 0.0079, Fig. 4c) and rightward in autoD2KD mice (p = 0.0029; Fig. 4c). Not only were the autoD2KD mice more exploratory compared to controls, they were among the most exploratory, even when compared to MSN-D2KD (p < 0.0001, Fig. 4c, Supplementary Fig. 1h-i) and double-D2KD (p < 0.0001). There were no differences between the double-D2KD and controls (p = 0.53, Fig. 4c, Supplementary Fig. 1h-i).

**Figure 4.**
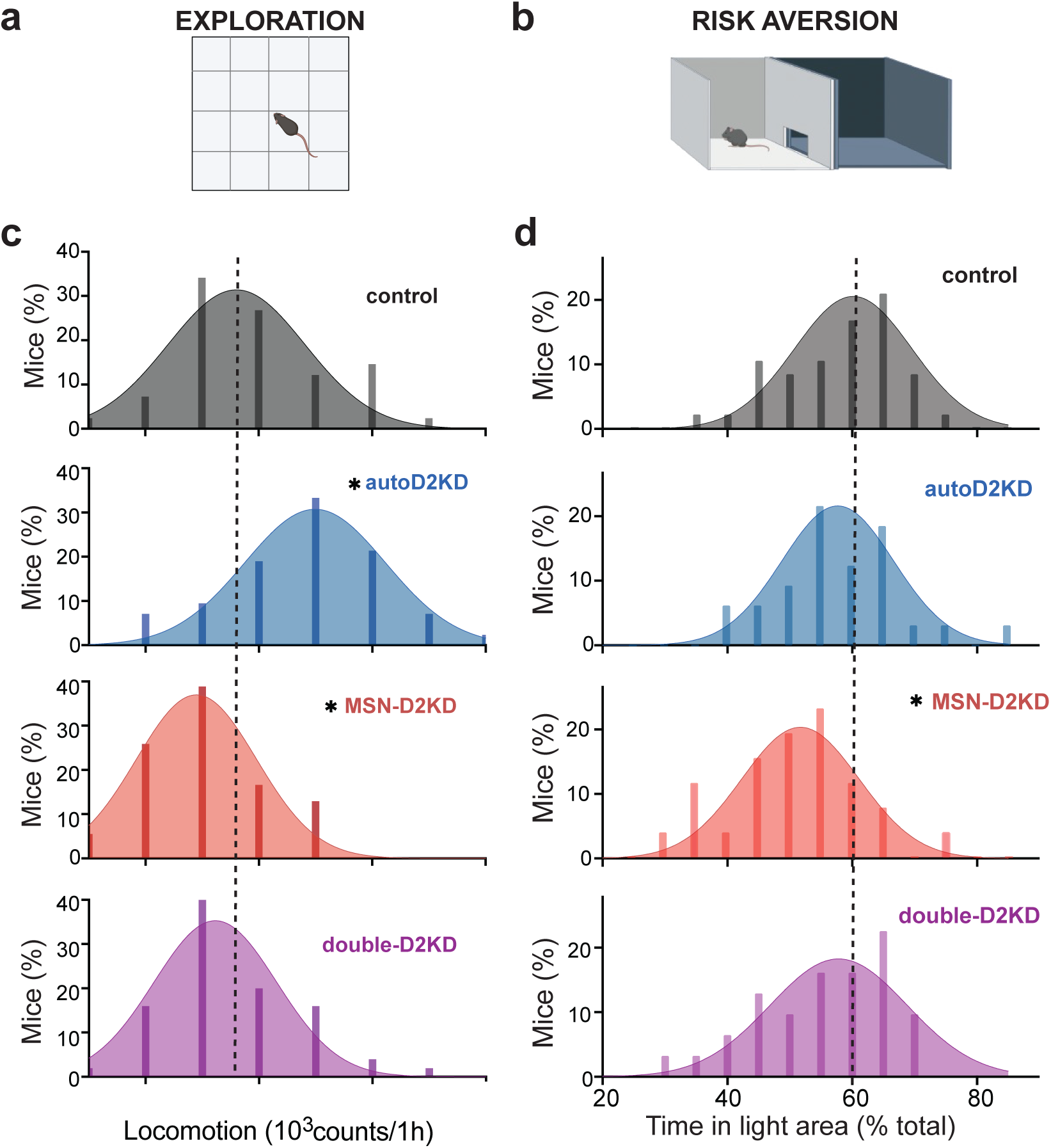
Cell-type–specific Drd2 haploinsufficiency produces divergent trait phenotypes. **a,** Novelty-driven exploration assessed in a novel open-field arena. **b,** Risk avoidance assessed in the light–dark box. **c,d,** Frequency distributions for novelty-induced locomotion (**c**) and time spent in the light compartment (**d**) for littermate mice of each genotype (n = 40 mice per genotype). Dashed lines indicate the mean of the control group. For all panels, *p ≤ 0.05 versus control.

To exclude global motor differences as a confounding factor, we measured baseline locomotion in the home cage across multiple days. All genotypes exhibited normal circadian rhythms, with higher activity during the dark phase (main effect of light phase: F(1,56) = 239, p < 0.0001; Supplementary Fig. 1e-f). A modest overall genotype effect did not reach significance (F(3,56) = 2.643, p = 0.058), and none of the knockdown genotypes differed from controls in total movement during either the light or dark phase (Supplementary Fig. 1). Together, these data indicate that reduced novelty-induced exploration in MSN-D2KD mice reflects altered exploratory drive rather than generalized hypoactivity.

### Risk avoidance behaviors and developmental modulation

Risk avoidance was assessed via two approach–avoidance assays: the light–dark box and the elevated zero maze (Fig. 4b,d, Supplementary Fig. 2). In the light–dark box, overall genotype differences emerged (one-way ANOVA, F(3,119)= 3.32, p = 0.022). MSN-D2KD mice spent less time in the illuminated (risk-associated) compartment than controls (post hoc, p = 0.01; Supplementary Fig. 2). Consistent with previous findings^27^, the distribution of time in the illuminated zone was shifted leftwards in MSN-D2KD mice relative to controls, indicating increased risk avoidance (Fig. 4b). This leftward shift was specific to D2 heteroreceptor knockdown and Cre recombinase expression alone did not reproduce the phenotype in Adora2a-Cre or Dat-ires-Cre control mice, or in mice carrying both Cre drivers (p = 0.52; Supplementary Fig. 2).

A robust age-dependent shift in risk-related behavior was observed in the elevated zero maze between mice aged 3 and 6 months. Across genotypes, younger mice spent more time in the open zones and made more open-zone entries than 6-month-old mice (two-way ANOVA, main effect of age: F(1,173) = 49.37, P < 0.0001; Supplementary Fig. 2; Table S1). Genotype also influenced performance (genotype: F(3,173) = 2.98, p = 0.033) without a genotype × age interaction (F(3,173) = 1.54, p = 0.21). Genotypic differences were minimal at 3 months but evident at 6 months, when MSN-D2KD and double-D2KD mice spent less time in open zones than controls (p = 0.053 and 0.014) and made fewer open-zone entries (p = 0.0016 and 0.018) (Supplementary Fig. 2; Table S1). Thus, though risk avoidance increased with age across all genotypes, reduced D2 signaling in MSNs was associated with greater avoidance, especially in older adulthood.

### Low autoreceptor function promotes cocaine desensitization

To characterize the behavioral response to acute and repeated cocaine administration, mice were habituated to the locomotor chambers and saline i.p. injections, then received cocaine (15 mg/kg, i.p.) once daily for 5 days (Fig. 5). During habituation to locomotor chambers, locomotion trended as expected for each genotype, with MSN-D2KD mice exhibiting lowest activity and autoD2KD mice the highest activity.

**Figure 5.**
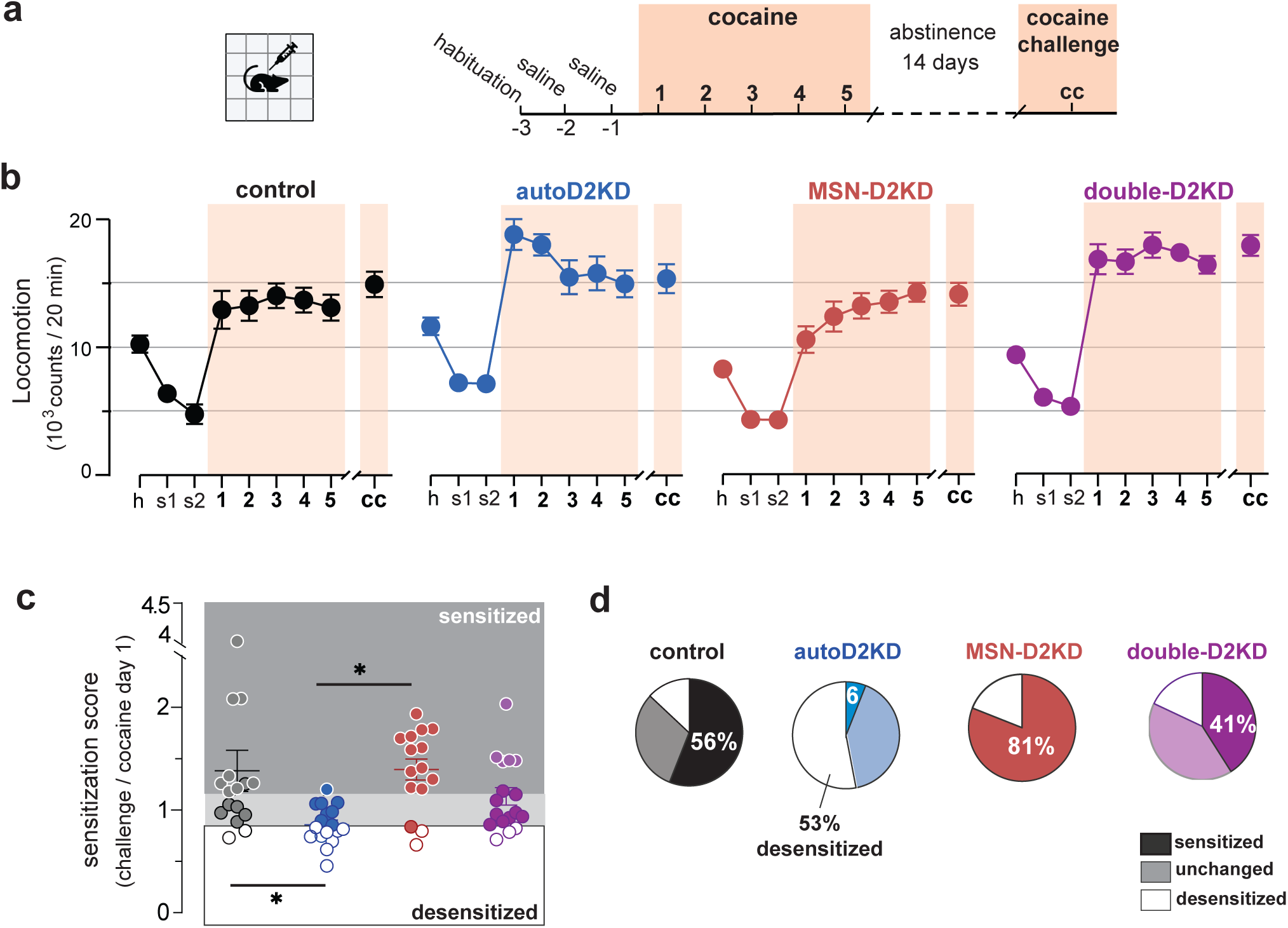
Acute cocaine stimulation and adaptations to repeated exposure. **a,** Experimental timeline. Mice were habituated to the activity chambers and saline injections (2 days), then received cocaine (15 mg/kg, i.p.) once daily for 5 consecutive days. Two weeks after the final injection, mice received a cocaine challenge dose (15 mg/kg, i.p.). **b**, Locomotor activity (20-min bins) during habituation (h), saline (s1, s2), cocaine days 1–5, and challenge (cc) across genotypes (n = 16 controls, 17 autoD2KD, 15 MSN-D2KD, 15 double-D2KD). Mean ± s.e.m. **c**, Sensitization score for each mouse (challenge/day 1 locomotion). Scores ≥ 1.15 were classified as sensitized and scores ≤ 0.85 as desensitized; intermediate values were classified as unchanged. Points show individual mice; horizontal lines indicate mean ± s.e.m. *p ≤ 0.05. **d**, Proportion of mice classified as sensitized, desensitized, or unchanged in each genotype.

Cocaine produced a robust increase in locomotion across groups (repeated-measures ANOVA, Cocaine: F(5.1,317.5) = 117.6, p < 0.0001) together with significant effects of genotype and a Genotype × Cocaine interaction (genotype: F(3,63) = 6.18, p = 0.001; genotype x cocaine: F(15.3,317.5) = 3.87, p < 0.0001, Fig. 5). autoD2KD mice exhibited an enhanced locomotor response to the first two cocaine injections (post hoc, day 1: p = 0.0123; p = 0.007 for day 2), but this effect diminished with continued exposure, such that responses were no longer different from controls by day 3. In contrast, MSN-D2KD mice showed a low initial response that increased across days, reaching levels comparable to autoD2KD mice by day 3. Double-D2KD mice displayed a sustained enhancement in cocaine-induced locomotion on days 3–5 (p = 0.02, 0.009 and 0.035, respectively) and were the only genotype to trend higher than controls during a cocaine challenge performed two weeks after the final exposure (p = 0.07).

To quantify longer-term adaptation, we computed a sensitization score as the ratio of the cocaine challenge response to the response on day 1. Mean sensitization scores were >1 for controls, MSN-D2KD and double-D2KD mice (1.4 ± 0.2, 1.4 ± 0.1 and 1.1 ± 0.08, respectively; Fig. 5c), whereas autoD2KD mice exhibited a significantly lower score (0.86 ± 0.05), reflecting a predominance of desensitization (one-way ANOVA: F(3,61) = 4.55, p = 0.0068; post-hoc Tukey’s: p = 0.014 and p = 0.015, respectively). Consistently, only 6% of autoD2KD mice sensitized (1 out of 17), compared to 56% of controls (9 out of 16), 81% of MSN-D2KD (13 out of 16) and 41% of double-D2KD mice (7 out of 17). Most autoD2KD mice were instead desensitized (53%), showing a smaller cocaine response at challenge than on day 1 (Supplementary Fig. 3). Thus, reduced D2 autoreceptor function was associated with heightened sensitivity to initial cocaine exposure followed by desensitization with repeated dosing, whereas reduced D2 signaling in striatal MSNs was associated with robust sensitization.

Cocaine reward learning was assessed using conditioned place preference (CPP). Mice were conditioned with distinct floor cues paired with cocaine (15 mg/kg) or saline on alternating sessions (Supplementary Fig. 3b) and were unbiased at baseline (genotype F(3,67)=0.123, p=0.95). Conditioning increased preference for the cocaine-paired floor across tests (main effect of test: F(1.9,123.4) = 45.14, p < 0.0001), with no effect of genotype (F(3,71) = 0.60, p = 0.62) and no Genotype × Test interaction (F(6,128) = 0.87, p = 0.52). Preference increased similarly across genotypes. Thus, CPP acquisition was intact despite Drd2 haploinsufficiency, whereas cocaine-evoked locomotor adaptations diverged by cell type.

### Acquisition of intravenous cocaine self-administration is intact across genotypes

Cocaine voluntary consumption and SUD–relevant phenotypes were assessed in an operant intravenous self-administration (IVSA) task that assayed (i) futile seeking during signaled drug unavailability, (ii) consumption under risk of punishment, (iii) under high-effort conditions, and (iv) craving-like seeking after short versus prolonged abstinence (1 day vs 5 weeks) (Fig. 6a-b).

**Figure 6.**
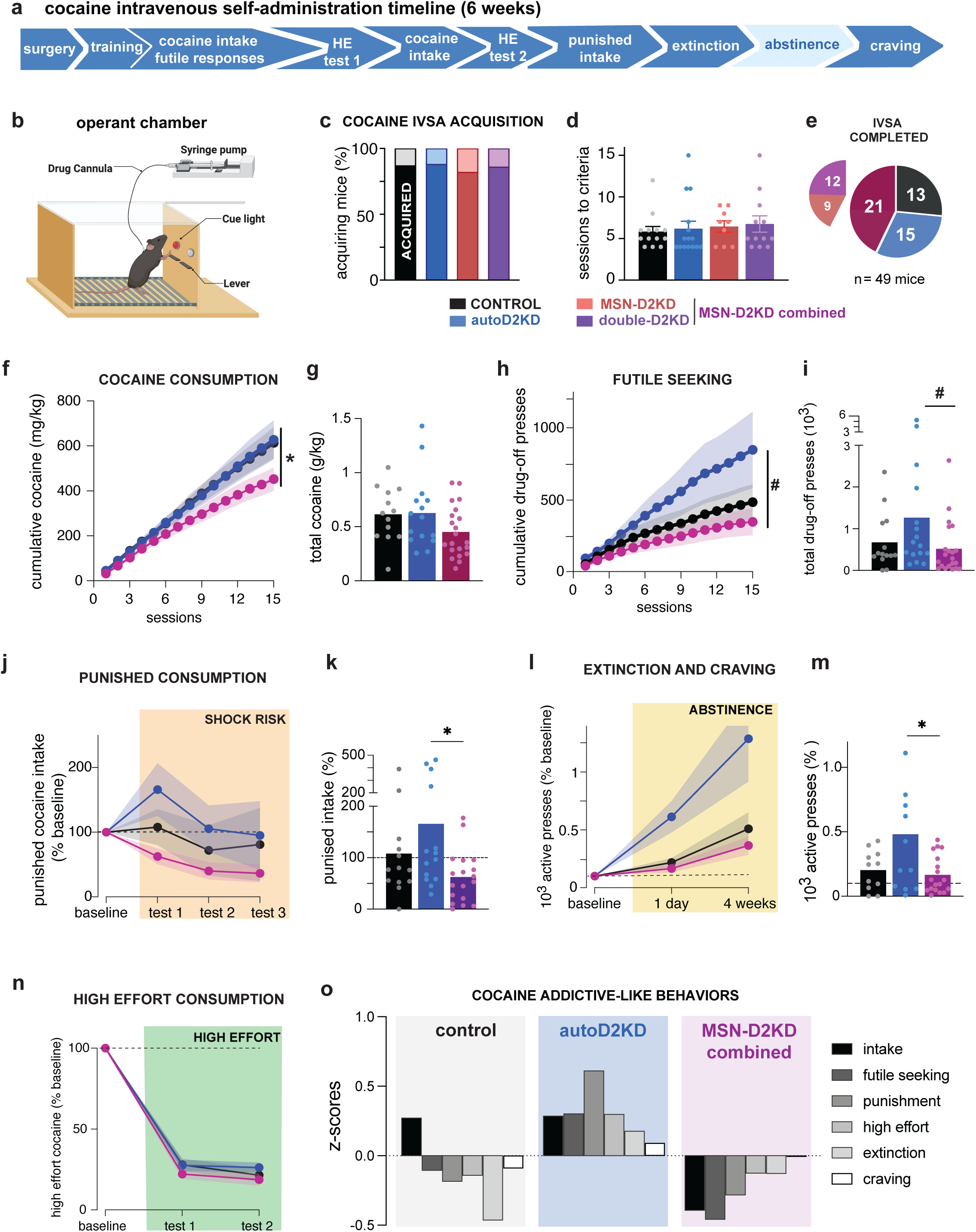
Operant cocaine self-administration reveals elevated addiction-like behavior in autoD2KD mice. **a,** Timeline of the intravenous self-administration (IVSA) protocol (∼6 weeks), including measures of futile seeking during signalled drug unavailability, punishment-resistant intake, high-effort consumption (HE), and cue/context-driven seeking after abstinence **b**, Schematic of the operant chamber showing active and inactive levers and the infusion pump for intravenous cocaine delivery. **c,** Proportion of littermate mice in each genotype that met acquisition criteria (dark shading) versus failed to acquire (light shading). **d,** Sessions required to reach acquisition criteria. Points denote individual mice; bars show mean ± s.e.m. **e,** Attrition across the protocol: number of littermate mice per genotype that completed the full IVSA schedule. For this figure, MSN-D2KD and double-D2KD mice are pooled and shown as “MSN-D2KD combined” groups; genotypes are shown separately in Supplementary Figs. 5–6. **f,h,** Across-session trajectories for cumulative cocaine intake (**f**) and futile responding during the drug-off period (**h**) over 16 IVSA sessions for controls (n = 13, black), autoD2KD (n = 15, blue) and heteroreceptor knockdown (n = 21, maroon). Symbols show group means; shaded bands indicate ± s.e.m. Genotype by Behavior interaction *p=0.003; #p=0.07 **g,i,k,m,** Summary plots for cumulative cocaine intake (**g**), cumulative futile responding (**i**), cocaine intake during punishment test 1 (**k**), and seeking on the first day of abstinence (**m**) for controls (black), autoD2KD (blue) and MSN-D2KD combined (maroon). Points denote individual mice; bars show mean. *p < 0.05; #p=0.07 post hoc after one way ANOVA **j,** Punishment-resistant intake across three tests, expressed as cocaine intake during punishment normalized to each mouse’s intake during unpunished sessions; one-third of earned infusions were paired with foot shock. **l,** Cue/context-driven seeking during abstinence ( day 1 extinction and week 4 craving), expressed as active lever presses during extinction tests normalized to each mouse’s responding during the final IVSA sessions. **n,** High-effort cocaine intake assessed with progressive-ratio tests (test 1 early; test 2 after extended IVSA), expressed relative to FR3 intake. For panels j, l, n, symbols show group means; shaded bands indicate ± s.e.m. **o**, Genotype mean z-scores (bar) for each cocaine-related measure. Individual z-scores were computed for each behavior using the cohort-wide mean and s.d. (collapsed across all four genotypes), then averaged within genotype. Main effect of genotype, p=0.002.

Mice were implanted with jugular catheters and trained to lever-press for intravenous cocaine under a fixed-ratio schedule (FR1, then FR3). A key control for these experiments is that all cohorts were littermates controlled across genotypes. Of 71 mice entering the protocol, 80% (57 mice) remained healthy with patent catheters through the end of operant training, and similar proportions across genotypes met acquisition criteria: 87% (n = 13/15) controls, 88% (15/17) autoD2KD, 82% (9/11) MSN-D2KD and 86% (12/14) double-D2KD (X^2^ = 0.24, p = 0.97; Fig. 6c). The number of sessions required to reach acquisition, defined by active-lever discrimination and cocaine intake (≥ 10 mg/kg/day), did not differ by genotypes (one-way ANOVA F(3,45) = 0.22, p = 0.88; Fig. 6d), indicating intact acquisition across groups.

### Severity of cocaine-related behaviors differs across cell-type-specific D2 knockdowns

During standard reinforcement conditions, cumulative cocaine consumption differed across genotypes over sessions, yielding a significant genotype × session interaction (F(28,644) = 1.9, p = 0.004; Figs. 6f-g, Supplementary Fig. 5). Intake trajectories overlapped for controls and autoD2KD mice, whereas MSN-D2KD and double-D2KD mice accumulated intake more slowly across sessions (Supplementary Fig. 5e). Consistent with this, pooling MSN-D2KD and double-D2KD (heteroreceptor knockdown group) revealed a trend toward lower cumulative consumption relative to autoD2KD mice (452 ± 237 mg/kg versus 627 ± 337 mg/kg; t = 1.8, p = 0.075). Across the remaining cocaine-related measures, MSN-D2KD and double-D2KD mice showed similar phenotypes (Supplementary Figs. 5-6). Accordingly, these groups were pooled for subsequent analyses and are shown as a combined heteroreceptor knockdown group (MSN-D2KD combined) in the main figures.

To probe inhibitory control over cocaine seeking, we quantified futile responding during a signaled drug-unavailable period inserted into each daily session (130 min access → 90 min drug-off → 130 min access). Cumulative futile responding showed a significant genotype × session interaction (F(28,672) = 2.05, p = 0.001; Fig. 6h-i). autoD2KD mice made more active lever presses during drug-off periods (1263 ± 375) than controls (579 ± 107) and the heteroreceptor knockdown group (518 ± 141) (one-way ANOVA, F(2,70) = 3.89, p = 0.025; post hoc p = 0.035; Fig. 6i, Supplementary Figs. 5-6).

We next assessed punishment-resistant intake by pairing a noxious foot shock with one-third of earned cocaine infusions over three daily tests. Punishment suppressed intake overall, but responses varied markedly across individuals (Supplementary Fig. 5; mouse effect: F(41,82) = 8.98, p < 0.0001). There were trends toward a main effect of genotype (F(2,43) = 2.84, p = 0.069) and a Genotype × Punishment interaction across the three tests (F(4,84) = 2.17, p = 0.071; Fig. 6j). Most control mice (62%) reduced consumption under punishment, whereas most autoD2KD mice (64%) maintained or increased intake. This reduced punishment sensitivity was selective to autoreceptor knockdown (one-way ANOVA, F(2,53) = 4.77, p = 0.012), and the heteroreceptor knockdown group showed robust suppression of intake under punishment (Wilcoxon test, p = 0.006, Fig. 6k).

Cocaine seeking during abstinence was assessed on the first day of extinction and after prolonged (4 weeks) abstinence to assess craving. Cocaine was replaced by saline while cocaine-associated cues were preserved. Across genotypes, response was elevated on the first extinction day relative to the final IVSA session (F(1.1,43.8) = 43.8, p = 0.0004), with a trend toward a genotype effect (F(2,38) = 2.83, p = 0.071, Fig. 6l). Extinction responding was highest in autoD2KD mice relative to the heteroreceptor knockdown group (one-way ANOVA, F(2,38) = 4.69, p = 0.015; post hoc p = 0.015 versus heteroreceptor knockdown; p = 0.065 versus controls, Fig. 6m).

Finally, increasing the response requirement (high-effort test) reduced cocaine consumption across pooled genotypes (main effect of test: F(2,86) = 651, p < 0.0001), with no effect of genotype (F(2,43) = 2.06, p = 0.139) (Fig. 6n). No differences were detected at either the early or late test (ps<0.13).

When cocaine intake and related measures were expressed as z-scores relative to the cohort mean, control mice showed a modestly positive mean for consumption but negative mean scores for the remaining behavioral measures (Fig. 5o). In contrast, autoD2KD mice exhibited positive z-scores across all measures (cocaine consumption, futile seeking, punishment-resistant intake, high-effort responding, and abstinence-related seeking) indicating broadly elevated cocaine-related behavior relative to the population. Conversely, the heteroreceptor knockdown group showed negative z-scores across measures, consistent with below-mean cocaine-related behaviors. A main effect of genotype was found (F92,46) = 5.9, p = 0.005).

### Intact sucrose preference and punishment sensitivity across genotypes

To assess whether genotype produced generalized differences in reward sensitivity, we measured operant sucrose self-administration in socially housed mice using the IntelliCage system, in which up to 16 same-sex mice per cage had operant access to water and 1% sucrose solution in separate corners (Supplementary Fig. 7). Sucrose preference relative to water was calculated per mouse per day and did not differ across genotypes (one-way ANOVA, F(3,56) = 1.21, p = 0.31; Supplementary Fig. 7). There was a trend toward higher sucrose preference in double-D2KD mice relative to controls (t(35) = 1.8, p = 0.0795), warranting follow-up.

We next paired sucrose access with a brief air-puff delivered on one-third of operant events. This punishment contingency suppressed sucrose consumption across all genotypes over three days, with no genotype differences on punishment day 1 (F(3,56) = 1.76, p = 0.17) or across the mean of punishment days 1–3 (F(3,56) = 1.83, p = 0.15). Notably, mice with reduced D2 autoreceptors did not differ from controls in their response to air-puff punishment, indicating that the punishment resistance observed during cocaine intake in autoD2KD mice does not generalize to aversive pairing during sucrose reinforcement.

### AutoD2KD mice are twice as likely to exhibit positive addiction-like composite scores

Although genotype differences in specific cocaine-related behaviors provide mechanistic insight, cocaine use disorder is clinically defined by aggregated severity across multiple behavioral domains. To mirror this framework, we performed an individual-level analysis in which cocaine-related measures obtained during voluntary cocaine self-administration were aggregated for each mouse (Fig. 7). For each animal, z-scores for cocaine intake, futile seeking, punishment-resistant intake, extinction responding, abstinence-related seeking/craving and high-effort responding were combined into a composite addiction-like score.

**Figure 7.**
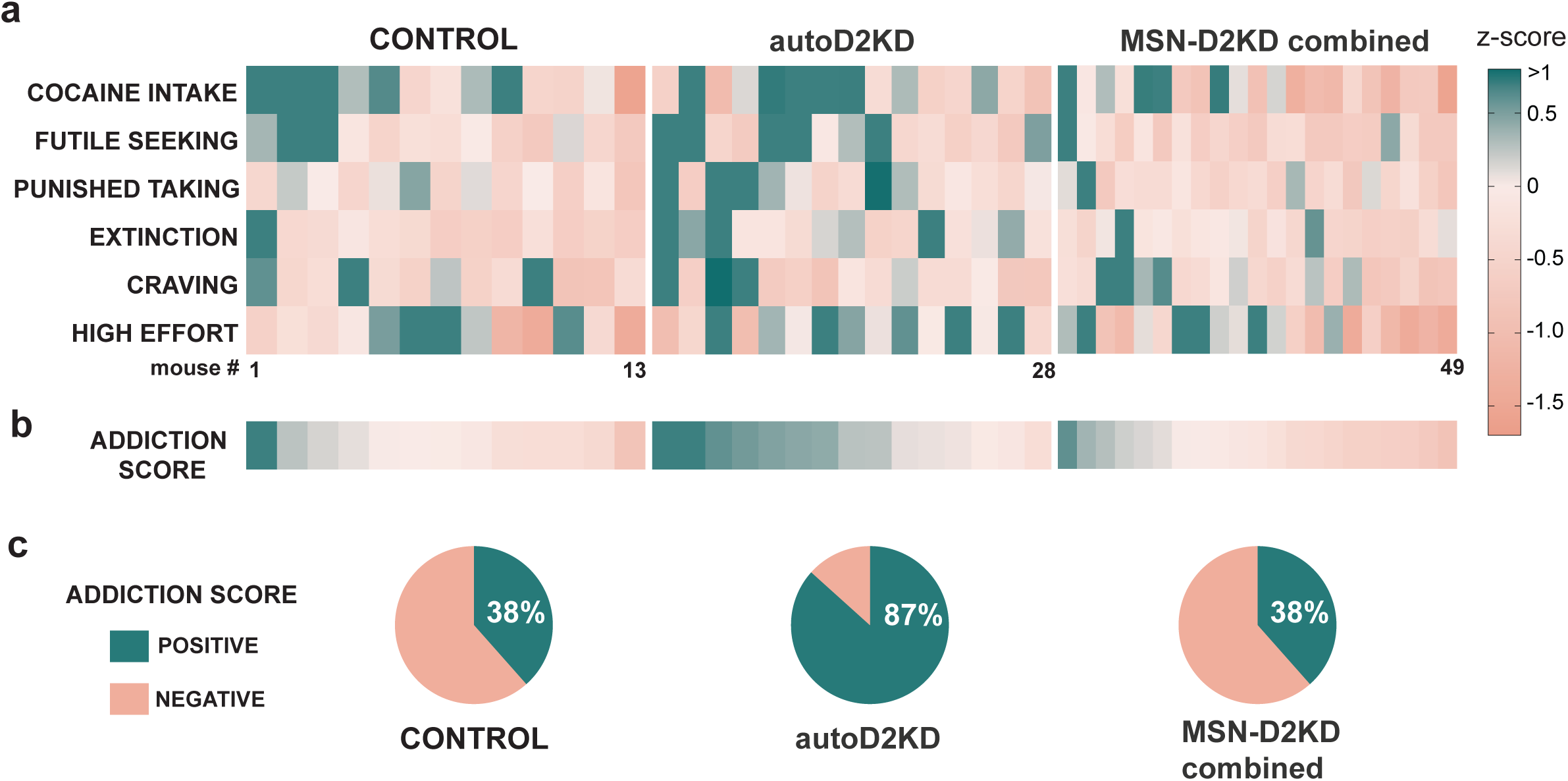
Autoreceptor haploinsufficiency increases the prevalence of addiction-like behavior. **a,** Heat map of individual mouse z-scores across the six cocaine-related behavioral measures (columns) for control, autoD2KD, and MSN-D2KD combined (MSN-D2KD+double-D2KD) mice. Main effect of genotype F (2, 264) = 7.624, p=0.0006. **b,** Composite addiction-like score for each mouse, calculated by integrating z-scores across measures (mean z-score per animal). For panels a and b, color scale indicates positive (green) and negative (orange) z-scores. One-way ANOVA genotype F (2, 46) = 6.872, p=0.002 **c,** Pie charts showing the proportion of mice in each genotype with positive (green) versus negative (orange) composite addiction-like scores. Genotype difference p=0.007 Fisher’s exact.

Using this metric, 38% of control mice exhibited a positive addiction-like score. A similar proportion was observed among mice with reduced D2 heteroreceptors in MSNs (38% pooled; 43% in MSN-D2KD; 33% in double-D2KD; Fig. 7b). In contrast, 87% of autoD2KD mice showed positive composite scores, corresponding to an approximately twofold increase in the prevalence of above-mean addiction-like behavior. This shift in proportions was significant (Fisher’s exact test, p = 0.007). Accordingly, autoD2KD mice exhibited higher addiction-like scores than both controls and mice with reduced D2 heteroreceptors (MSN-D2KD and double-D2KD).

## DISCUSSION

This study dissociates the behavioral and circuit consequences of reduced presynaptic D2 autoreceptors on dopamine axons from those of reduced postsynaptic D2 heteroreceptors on striatal medium spiny neurons. This distinction addresses a central challenge in understanding stimulant use disorder: reduced striatal D2/3 receptor availability is a robust PET correlate, yet the PET signal aggregates multiple receptors and compartments with distinct functions^33,34^. Using cell-type–specific Drd2 haploinsufficiency, we show that partial autoreceptor reduction—comparable in magnitude to clinically reported reductions in striatal D2/3 availability in stimulant use disorder—weakens presynaptic negative feedback control of dopamine release, increases phasic-to-tonic dopamine gain, and prolongs cocaine-evoked dopamine elevations. These physiological changes were accompanied by heightened novelty exploration, increased sensitivity to initial cocaine, and paradoxical desensitization with repeated exposure. In operant cocaine self-administration, D2 autoreceptor knockdown selectively increased SUD-like phenotypes across domains of inhibitory control and compulsive intake, whereas D2 heteroreceptor knockdown produced a contrasting trait profile and distinct dopamine receptor remodeling. Together, these data identify D2 autoreceptors as a key determinant of stimulant vulnerability and suggest that D1 receptor adaptations provide a readout of compartment-specific dopaminergic perturbations with relevance for risk stratification.

### Autoreceptor hypofunction selectively amplifies phasic dopamine and prolongs cocaine-evoked dopamine elevations

Mechanistic insight into vulnerability emerged from direct measurements of evoked dopamine release and its regulation by D2/3 agonists and cocaine. Exogenous D2/3 agonism produced weaker inhibition of dopamine release in autoD2KD mice, providing functional validation of reduced autoreceptor control. Under physiological conditions, however, autoreceptor feedback is preferentially recruited when extracellular dopamine rises—during high-frequency firing, high release probability, reduced reuptake, or combinations thereof. Consistent with this, tonic dopamine release evoked by low-frequency stimulation was unchanged in autoD2KD mice, whereas high-frequency stimulation revealed larger dopamine signals when expressed relative to tonic release, indicating enhanced phasic-to-tonic gain. This physiological selectivity paralleled locomotor behavior: genotype differences were modest in a familiar environment but amplified by novelty, with autoD2KD mice showing heightened exploration.

Cocaine increased dopamine signal amplitude and slowed decay in both genotypes, consistent with dopamine transporter blockade. In controls, this potentiation waned over time, whereas in autoD2KD mice it persisted, yielding prolonged cocaine-evoked dopamine elevations—consistent with weakened autoreceptor-mediated feedback. Behaviorally, autoD2KD mice were hypersensitive to initial cocaine yet rapidly desensitized with repeated exposure. The mechanisms underlying this paradoxical adaptation remain to be defined, but our receptor-binding data suggest a plausible contribution of reduced D1 signaling. In NAc D2-MSNs, repeated cocaine reduces postsynaptic D2 receptor sensitivity through a D1-dependent process, as pretreatment with the D1R antagonist SCH23390 blocks it^28^. We found coordinated downregulation of D1-like binding in autoD2KD mice, which may bias behavioral responses toward blunted sensitization and desensitization despite prolonged dopamine elevations. In this view, reduced D1-like binding in autoD2KD mice could dampen D1-dependent cocaine plasticity and bias toward desensitization, whereas the elevated D1:D2/3 ratio in MSN-D2KD mice may facilitate D1-dependent adaptations that support sensitization.

Indeed, desensitization was selective to mice with reduced autoreceptor function: mice with reduced D2 heteroreceptors showed blunted acute stimulation and robust sensitization. The dissociation observed here—elevated addiction-like behavior in autoD2KD mice despite desensitization, and robust sensitization in heteroreceptor knockdown mice without elevated addiction-like behavior—underscores that locomotor sensitization and compulsive cocaine seeking/taking can be mechanistically separable.

### D1 remodeling and the D1:D2/3 balance distinguish D2 receptor compartments

A translationally relevant finding is that D1-like receptor adaptations depended on the D2 compartment perturbed. *A priori,* reducing D2 receptors might be expected to increase the D1:D2/3 binding ratio irrespective of cell type. Instead, the ratio increased only when D2 heteroreceptors were reduced in MSNs—either alone or in combination with autoreceptor knockdown. By contrast, autoreceptor knockdown reduced D2/3 binding without increasing the ratio because it was accompanied by a coordinated reduction in D1-like binding. Thus, D1-like binding, together with the D1:D2/3 ratio, provides a potential strategy to infer the compartmental basis of “low D2/3” signals.

Double-D2KD mice exhibited a regionally mixed phenotype: the D1:D2/3 ratio was elevated in dorsal striatum but near control in nucleus accumbens. In accumbens, ratio normalization appeared to reflect a smaller reduction in D2/3 binding coupled to lower D1-like binding than in MSN-D2KD mice. An additional possibility is compensatory changes in D3 receptors, which contribute to raclopride-class binding in ventral striatum and could partially buffer apparent D2/3 losses.^35,36^

### Traits, development, and vulnerability: linking receptor balance to risk avoidance and novelty seeking

Across genotypes, the D1:D2/3 ratio behaved as a compact readout of dopaminergic “balance” with implications for trait expression. Mice with elevated dorsal striatal D1:D2/3 ratios (MSN-D2KD and double-D2KD) exhibited increased risk avoidance^27^. This phenotype was developmentally modulated: risk avoidance increased with age across genotypes, and the age-related shift was most pronounced in mice with elevated D1:D2/3 ratios, suggesting a gene-by-development interaction in the maturation of frontostriatal circuits.

In contrast, autoD2KD mice exhibited a distinct configuration, lower D2/3 binding accompanied by reduced D1-like binding, which was associated with heightened novelty exploration and elevated cocaine SUD–like behavior. These findings align with prior work linking midbrain dopamine receptors to novelty seeking and stimulant vulnerability^30,37–39^. Human imaging studies further suggest that reduced midbrain autoreceptor function and impaired dopaminergic regulation contribute to compulsive disorders^34^. More recent clinical data showing that caudate D1:D2/3 ratios positively correlate with attentional capacity and negatively with methylphenidate improvements of attention converge with our results^40^. Both these studies support a model in which weak D2 autoreceptor control biases individuals toward greater benefit from stimulants and this normalizing effect of stimulants may potentially increase vulnerability to repeated use^34,38,39,41^. In this sense, the normalization of hyper-exploration by repeated cocaine in autoD2KD mice is consistent with a self-medication framework, although directly testing this hypothesis will require assays of performance, internal-state relief, and reinforcement contingencies that isolate “normalization” as the reinforcer.

This work suggests that a self-medication framework may apply differently across substances depending on the compartment-specific nature of reduced D2 signaling. In a complementary alcohol study, MSN-D2KD mice—which exhibit an elevated D1:D2/3 ratio and a risk-avoidant trait—show stronger anxiolytic response to alcohol and increased compulsive-like alcohol behaviors, consistent with the possibility that alcohol normalizes their anxiety state^27^. By contrast, autoD2KD mice may be preferentially vulnerable to compulsive stimulant use in part because cocaine normalizes a hyper-exploratory baseline phenotype. More broadly, these findings raise the possibility that “low D2/3” PET signals observed across SUDs could arise from distinct combinations of presynaptic autoreceptor and postsynaptic heteroreceptor alterations, with different behavioral consequences and treatment implications. Testing this hypothesis will require a biomarker that can resolve these compartments and cross-drug comparisons within the same experimental framework^42^.

### Translational scale and specificity

A longstanding concern in translational addiction research is whether animal perturbations approximate the magnitude of dopaminergic alterations in humans. PET studies in cocaine dependence typically report modest (∼15–17%) reductions in striatal D2/3 availability^3^, and low D2/3 availability in healthy individuals predicts greater subjective and neural response to methylphenidate^2,4,13^. In this context, the relatively small reduction in D2/3 binding in autoD2KD mice supports its relevance as a model of clinically observed “low D2/3” states. Notably, the strongest cocaine SUD–like phenotypes for cocaine emerged in the genotype most closely aligned with clinical effect sizes, arguing that modest reductions in autoreceptor function are sufficient to bias traits and increase stimulant vulnerability.

Behavioral specificity further supports this interpretation. We did not detect genotype differences in sucrose preference or punishment sensitivity during sucrose reinforcement, arguing against generalized reward/punishment deficits. CPP for cocaine was also intact across genotypes, suggesting that vulnerability in autoD2KD mice is not driven by gross enhancement of cocaine’s conditioned reward learning. Together, these patterns are consistent with the possibility that dependence-related behaviors can arise from state normalization and altered control processes rather than from uniformly enhanced reward learning^39^.

### Limitations and technical considerations

Several limitations constrain generalization. First, these findings were obtained using cocaine, and dopaminergic mechanisms contributing to vulnerability may differ across drug classes. Further, among recreational users, concurrent use of other substances to minimize or counteract stimulant side effects likely altered drug responses, further constraining the generalization. Second, Drd2 haploinsufficiency was present from early development which argues it is a preexisting causal factor; however, adult-onset manipulations will be necessary to dissociate developmental from adult circuit contributions. Third, although the study included both sexes and did not identify robust sex differences across the measures tested, it might be unpowered to exclude subtle sex-by-genotype interactions. Fourth, the dopamine physiology experiments used Dat-ires-Cre+ controls to match the genetic background of autoD2KD mice, given evidence for modest DAT expression/kinetic differences in this line; future work should test generalization across alternative genetic strategies (e.g., En1-Cre–based autoreceptor targeting) and across independent cohorts.

## Conclusion

Taken together, these results provide a cell type-specific interpretation of reduced striatal D2/3 signaling that resolves a key ambiguity in results observed from human studies. Partial loss of D2 autoreceptor function selectively amplifies phasic dopamine signaling and prolongs cocaine-evoked dopamine elevations, biasing baseline traits and increasing the probability of cocaine SUD–like behaviors across DSM-aligned domains. In parallel, D1 remodeling—and the resulting D1:D2/3 balance—distinguishes heteroreceptor versus autoreceptor perturbations and links dopaminergic “balance” to trait axes such as risk avoidance. These findings nominate D2 autoreceptors as a mechanistic determinant of stimulant vulnerability and motivate development of biomarker strategies that combine D2/3 measures with D1 metrics to improve risk stratification and mechanism-guided intervention.

## Supporting information

STATISTIC TABLE

## METHODS

### Animals

Experiments were performed in accordance with guidelines from the National Institute on Alcohol Abuse and Alcoholism’s Animal Care and Use Committee. Mice homozygotes for Drd2^loxP/loxP^ (B6.129S4(FVB)-Drd2<tm1.1Mrub>/J; IMSR_JAX:020631)^24^ and *Adora2a-Cre^+/-^*(B6.FVB(Cg)-Tg(*Adora2a-Cre*)KG139Gsat/Mmucd, GENSAT, 036158-UCD) (Gerfen et al 2013); were crossed with mice heterozygous DAT^IRES-CRE^ (B6.SJL-Slc6a3<tm1.1(cre)Bkmn>/J; IMSR_JAX:006660) (Backman et al. 2006). The genotypes of the pups generated were: autoD2KD (Drd2^loxP/w^; DAT^IRES-Cre^); MSN-D2KD (Drd2^loxP/w^;Adora2A-Cre*^+/^*-); double-D2KD (Drd2^loxP/w^;Adora2A-Cre*^+/^*-;DAT^IRES-Cre^); and controls mice (Drd2^loxP/w^). For all experiments, mice of both sexes were used and counterbalanced; mice were housed with food and water available *ad libitum*. Each experiment consisted of separate cohorts of mice, except for locomotor response to D1-like and D2/3 agonists that tested the same mice. With the exception of cocaine IVSA experiments, mice were group-housed under a 12 h light cycle (6:30 ON/18:30 OFF) and behavioral testing took place during the light phase of the cycle.

### Quantitative polymerase chain reaction

Mice (males only, 14-31 weeks old) were anesthetized with isoflurane and decapitated. The brain was removed and brain regions (cortex, dorsal striatum, nucleus accumbens, amygdala, hippocampus, and ventral tegmental area) were dissected on ice, homogenized, and RNA was purified using RNeasy Plus Mini kit (Qiagen). cDNA was synthesized using iScript Reverse Transcription Supermix (Biorad). Relative dopamine D2 receptor (Oa04895884_m1 TaqMan, Catalog #: 4351372) and beta-actin probe (Mm01205647_g1 TaqMan, Catalog #: 4331182) mRNA expression were determined with TaqMan Gene Expression Assays (Life Technologies) using a StepOnePlus Real-Time PCR system (Applied Biosystems). Relative *Drd2* expression was calculated using the ΔΔCt method.

### Autoradiography

mice (both sexes, 10-20 weeks old) were anesthetized with isoflurane and brain removed quickly. Whole brains were flash-frozen in 2-methylbutane and stored at -80°C until sectioning (20 μm) on a Cryostat (CryoStar NX-50, Epredia) where slices were thaw mounted onto ethanol-washed glass slides. Slides were preincubated (10-15 minutes, room temperature [RT]) in washing buffer (50 mM Tris-HCl, pH 7.4), transferred to incubation buffer (120 minutes, RT; 50 mM Tris-HCl, pH 7.4, with 120 mM NaCl and 10 mM MgCl; for 60 minutes with 1 mM MgCl2, 5 mM KCl, 2 mM CaCl2]) containing either [3H]raclopride (4 nM, 80.8 Ci/mmol, PerkinElmer, or 81.8 Ci/mmol, Revvity) or [3H]SCH23390 (2.5 nM, 83.6 Ci/mmol, PerkinElmer, or 83.9 Ci/mmol, Revvity), with Ketanserin Tartrate (40 nM, Tocris) to block off-target serotonergic binding by SCH-23390. Non-specific binding of [3H]raclopride or [3H]SCH23390 was determined by incubating a subset of slides in the same conditions in the presence of either butaclamol (10 μM, Tocris) or SCH-23390 (10 μM, Tocris), respectively. Slides were then washed 2 x 15 seconds in ice-cold Tris-HCl buffer (50mM) and dipped in ice-cold distilled water to remove salts. The slides were dried overnight and apposed to a BAS-TR2025 Storage Phosphor Screen (Fujifilm) along with a Carbon-14 Standards slide (American Radiolabeled Chemicals Inc.) inside a Hypercassette (Amersham Biosciences) for 12 days and then imaged using a Typhoon biomolecular imager (Cytiva). Images were calibrated and analyzed using ImageJ 1.51j8 (NIH) or Multigauge software (GE Healthcare). Regions of interest (ROI; either 8-12 or 24-30 ROI per region and animal) were drawn freehand based on neuroanatomical landmarks and quantified by densitometry. Specific binding was calculated by subtracting non-specific binding (nCi/g) from each ROI, and percent binding was determined by normalizing these values to the mean specific binding in the corresponding striatal subregions of control mice.

### Locomotor activity and approach-avoidance assays

#### Novel open-field exploration

Mice (5–24 weeks old; both sexes) were transferred to the testing room and acclimated for 30–60 min before placement in clear polycarbonate chambers (20 cm H × 17 cm W × 28 cm D) equipped with infrared photobeam detectors (Columbus Instruments). Locomotor activity was recorded for 60 min. Unless otherwise stated, all locomotor assays used this beam-break system.

#### Home-cage-like locomotion

Mice (both sexes) were placed individually in the same photobeam chambers (20 cm H × 17 cm W × 28 cm D) fitted with food and water, and activity was recorded continuously for 48 h across light and dark phases. Total beam breaks were summed in 12-h bins and averaged across the two days to yield mean light-phase and dark-phase activity per mouse. Data for control and MSN-D2KD mice were previously reported^27^ and were reanalyzed here alongside autoD2KD and double-D2KD groups.

#### D1-like and D2/3 agonist induced locomotion

Testing was conducted during the light phase and all intraperitoneal drug administration was delivered at 10 ml kg^−1^ body weight. To minimize novelty-induced locomotion, baseline activity was recorded for 60 min before injection^43^, followed by 120 min of post-injection recording. Mice were habituated to the chambers on day 1 and received saline (i.p.) on days 2–3. SKF-81297 hydrobromide (Tocris, CAS # 67287-39-2) was then tested using escalating doses (2.5, 5.0 and 7.5 mg kg^–1, i.p.), with 2–3 days between doses. After a 2-week washout period, Quinelorane hydrochloride (RD Systems, CAS# 97548-97-5) was tested using the same schedule (two saline days followed by escalating doses of 0.005, 0.01 mg kg^–1, i.p.).

#### Light-Dark Box

The apparatus was a polycarbonate chamber (44.5 × 45.5 × 39 cm) divided into an illuminated compartment (∼260 lux) and a covered dark compartment, connected by a small opening. Mice (box sexes, 12-14 weeks old) were placed in the center of the light compartment and allowed to explore for 10 min. Behavior was recorded from above (Panasonic WV-CP304) and analyzed in TopScan (CleverSys) to quantify time spent in each compartment and the number of transitions. Automated scoring of time in the light zone was manually verified for accuracy.

#### Elevated Zero Maze

The elevated zero maze consisted of a circular runway (50 cm inner diameter; 5 cm lane width) elevated 50 cm above the floor, with two open and two enclosed quadrants (wall height 15 cm). Mice (box sexes) were tested at 12-14 weeks old (3 months) and 24-27 weeks old (6 months) were placed in an open quadrant (∼110 lux) and allowed to explore for 10 min. Behavior was recorded from above (Basler acA1300-60gm GigE) and analyzed in EthoVision XT 17 (Noldus) to quantify time spent in open quadrants. Closed-quadrant position tracking was unreliable in most recordings, precluding distance-based measures. Mice that fell from the maze were excluded.

### Evoked Dopamine Signals

Brain slice preparation and dopamine recordings were performed as previously described^44^. Briefly, mice (11–26 weeks; both sexes) were anesthetized with isoflurane and decapitated, and brains were rapidly removed into ice-cold sucrose-substituted artificial cerebrospinal fluid (ssACSF) containing (in mM): 90 sucrose, 80 NaCl, 24 NaHCO₃, 10 glucose, 3.5 KCl, 1.25 NaH₂PO₄, 4.5 MgCl₂ and 0.5 CaCl₂, saturated with 95% O₂/5% CO₂. Coronal slices (240 µm) were cut on a vibratome (Leica VT1200S) and incubated for 20–30 min at ∼32 °C in ACSF containing (in mM): 124 NaCl, 26.2 NaHCO₃, 20 glucose, 2.5 KCl, 1 NaH₂PO₄, 2.5 CaCl₂, 1.3 MgCl₂ and 0.4 ascorbic acid (95% O₂/5% CO₂), then held at room temperature until recording. For recordings, slices were transferred to a submerged chamber and perfused with warmed ACSF (32 °C; 2 ml min⁻¹) using an in-line heater (Harvard Apparatus). *Fast-scan cyclic voltammetry* (FSCV): was performed in the nucleus accumbens and dorsal striatum. AutoD2KD mice (Drd2^loxP/wt^; DAT^IRES^ ^Cre^) and DAT^IRES^ ^Cre^ (IMSR_JAX:006660) controls were used for most experiments. For optogenetic stimulation experiments, these lines were crossed to Ai32 (IMSR_JAX:024109) to express channelrhodopsin-2 in dopamine axons. Carbon-fiber microelectrodes were fabricated from a cylindrical carbon fiber (7 µm diameter; ∼150 µm exposed) inserted into a glass capillary and backfilled with 3 M KCl. Electrodes were conditioned by applying an 8-ms triangular waveform (−0.4 to +1.2 to −0.4 V versus Ag/AgCl; 400 V s⁻¹) every 15 ms before recording. During recordings, the electrode was held at −0.4 V and the same waveform was applied every 100 ms. The electrode was inserted diagonally (∼22°) to increase tissue contact and minimize downward force. For electrical stimulation, a glass pipette filled with ACSF was positioned near the electrode tip and a 0.2-ms rectangular pulse was delivered every 1–2 min (SIU91A, Cygnus Technologies). All electrical-evoked recordings were performed in the presence of the nicotinic receptor antagonist DHβE (Dihydro-β-erythroidine hydrobromide, Tocis Cat. #2349, at 1 µM) to suppress cholinergic contributions to dopamine release. For optogenetic stimulation, 473-nm LED light (CoolLED pE-800) pulses (14.8 mW, 0.2–5 ms duration) were delivered through the objective (40×, NA=0.8 Olympus). (-)-Quinpirole hydrochloride (30 nM, CAS # 85798-08-9) and cocaine HCl (3 µM, CAS# 53-21-4) were prepared fresh in ACSF. Signals were acquired with a retrofit headstage (CB-7B/EC with 5 MΩ resistor) and a MultiClamp 700B amplifier (Molecular Devices), low-pass filtered at 3 kHz, and digitized at 100 kHz (NI USB-6229 BNC, National Instruments). Data acquisition and analysis used custom VIGOR routines in Igor Pro (WaveMetrics) with mafPC procedures (courtesy of M.A. Xu-Friedman).

### Acute and repeated cocaine-induced locomotion

Locomotor activity was measured for 60 min using infrared beam-break detectors (Columbus Instruments). Mice (6–11 weeks old; both sexes) were habituated to the locomotor chambers on day 1 and received saline injections (i.p.) on days 2–3. Mice then received cocaine (15 mg/kg, i.p.) once daily on days 4–8, and locomotion was recorded for 60 min after each injection. Two weeks after the final cocaine injection, mice received a cocaine challenge injection (15 mg/kg, i.p.) and were again monitored for 60 min. A sensitization index was calculated for each mouse as the ratio of locomotor activity on the challenge day to activity after the first cocaine injection. Mice were classified as sensitized if challenge-day locomotion increased by ≥15% relative to the first cocaine exposure, and as desensitized if challenge-day locomotion decreased by ≥15%.

### Cocaine conditioned place preference

Conditioned place preference (CPP) was conducted in unbiased, two-compartment clear polycarbonate chambers (33 × 54 × 20 cm; H × W × D; Applying Reason and Technology LLC) housed in sound- and light-attenuating cabinets with ventilation. Position was tracked by infrared video (Q-See) and locomotor activity by infrared photobeams (Columbus Instruments). Chambers were not illuminated during conditioning or testing. Compartments were distinguished by tactile floor cues: stainless-steel grid (1.9-mm woven rods) or perforated “hole” floors (4.7-mm holes, 1.6 mm spacing). Mice (5–23 weeks old; both sexes) were randomized to cocaine paired with grid (Grid+) or hole (Hole+) floors, and drug order on the first conditioning day was counterbalanced. Data were analyzed in EthoVision XT9 (Noldus).

Mice underwent a saline pre-test (30 min), eight conditioning sessions (30 min; one per day; alternating cocaine 15 mg/kg, i.p., and saline), and two saline preference tests (30 min) after the 4th and 8th conditioning sessions. During preference tests, mice had access to both compartments. CPP was quantified as time spent on the grid floor; conditioning was inferred from a difference between Grid+ and Hole+ groups in grid-floor time^45^.

### Operant intravenous cocaine self-administration (IVSA)

Cocaine IVSA and catheterization procedures were performed as previously described (Bock et al., 2013; Holroyd et al., 2015). Mice (both sexes; 8–10 weeks at surgery) were group-housed on a reversed light cycle (lights on 21:30–09:30) for ≥1 week before experiments, with food and water available ad libitum. Under isoflurane anesthesia, a chronic indwelling catheter (Instech CP20PU-MJV2010; cut to length) connected to a vascular access button (Instech VABM1VB/25) was implanted in the right jugular vein. Mice recovered for 5–7 d and received antibiotic prophylaxis in drinking water (sulfamethoxazole/trimethoprim; SMZ–TMP, NDC 70954-258-10; 7.5 ml of 200 mg/40 mg per 5 ml stock per 260 ml water). Mice were singly housed after surgery for the duration of IVSA. Catheters were maintained with daily saline flushes. Patency was verified before IVSA and every 10 d thereafter using a brief anesthetic test (ketamine 90 mg/kg and xylazine 10 mg/kg in saline); mice failing patency were excluded. Sessions were conducted during the dark phase in operant chambers (Med Associates) equipped with two levers, cue lights, a house light, and a syringe pump. Active-lever responses delivered an intravenous cocaine infusion (1 mg/kg per infusion over 2–4 s, adjusted to body weight) and were paired with illumination of the cue light above the active lever; the cue light was extinguished during infusion. Inactive-lever responses were recorded but had no programmed consequences. The house light signaled session state (off at session onset; on at session end and during drug-unavailable periods).

### IVSA training, testing and behavioral measures

#### Training and acquisition

Mice were trained in daily 4.5 h sessions with continuous access to cocaine (1 mg/kg per infusion) under a fixed-ratio 1 (FR1) schedule. Mice advanced to FR3 after meeting acquisition criteria: (i) active:inactive lever-press ratio ≥2.5 and (ii) cocaine intake >10 mg/kg/d in each of the final two training sessions.

#### Self-administration sessions (FR3)

Daily self-administration sessions comprised two drug-available (“drug-on”) epochs (2.25 h each) separated by a drug-unavailable (“drug-off”) epoch (1.5 h). During drug-off, the house light was illuminated and the active-lever cue light was extinguished; lever presses were recorded as futile responses but had no programmed consequences.

#### High-effort intake (progressive ratio)

Motivation for cocaine was assessed with two progressive-ratio (PR) tests: PR1 after six FR3 sessions and PR2 after an additional ten FR3 sessions. Response requirements increased according to the Richardson and Roberts (1996) function (ratio = 5e^(0.2 × infusion number) − 5)^46^. Each PR session lasted 5 h or ended 1 h after the last earned infusion. High-effort intake was quantified as rewards earned during PR normalized to each mouse’s mean rewards during the final two FR3 sessions. After PR test 2, mice completed three FR3 sessions to re-establish baseline intake.

#### Punishment risk

Punishment-resistant intake was assessed in three consecutive sessions identical to FR3 self-administration, except every third earned infusion was paired with a foot shock (500 ms, 0.25 mA) delivered through the grid floor using an aversive stimulator/scrambler (Med Associates). After punishment testing, mice completed two FR3 sessions under non-punished conditions.

#### Seeking and craving during abstinence

Cue/context-driven seeking was assessed under extinction conditions in two 60-min tests: (i) 24 h after the final FR3 session (early abstinence; seeking) and (ii) after 4 weeks of forced abstinence (late abstinence; craving). During both tests, cocaine was unavailable and responses on either lever had no programmed consequences, while the operant context and cues were maintained. During forced abstinence, mice remained in their home cages with food and water ad libitum.

#### Data acquisition and quantification

Operant data were collected using Med-PC IV and analyzed in Prism (GraphPad) and COBAI (Igor Pro; WaveMetrics)^31,47^. Of 146 mice undergoing catheter surgery, 78 (54%) were patent and healthy at the start of IVSA; 63% of these completed the 5-week protocol and were included in analyses. Futile responding was quantified as active-lever presses during drug-off epochs and expressed cumulative. Seeking and craving were quantified as active-lever presses during extinction tests normalized to active-lever responding during the final two FR3 sessions. High-effort intake was quantified as rewards earned during PR normalized to mean rewards during the final two FR3 sessions. For each IVSA measure, z-scores were computed using the cohort-wide mean and s.d. across all mice completing the protocol (all four genotypes). To reduce dependence of futile seeking on overall intake, futile seeking was z-scored from a normalized metric (active presses during drug-off divided by active presses during drug-on for each mouse). Independence among behavioral measures was evaluated using a correlation matrix (Supplementary Fig. 4b). A composite addiction-like score was calculated for each mouse as the mean of its z-scores across cocaine intake, futile seeking, punished intake, high-effort intake, seeking, and craving.

#### Sucrose drinking and punishment sensitivity (IntelliCage)

Operant sucrose drinking and sensitivity to an aversive outcome were assessed in socially housed mice using the IntelliCage system (TSE Systems). Mice (both sexes) were habituated to IntelliCage chambers from 4–6 weeks of age (up to 16 mice per sex per cage) with unrestricted access to water in all four corners. At 7–9 weeks, mice were briefly anesthetized and implanted subcutaneously with an RFID transponder (dorsocervical region) for individual tracking. Each corner contained an RFID antenna and two computer-controlled doors regulating access to bottles; the system logged visit timestamps, nose pokes and licks. Data were inspected in TSE Analyzer and processed in R. *Water training:* Mice were trained to obtain water under an FR1 contingency (door opening for 5 s). Each reward required a new corner visit. After reaching ≥60% successful trials, mice advanced automatically to FR3 for the remainder of the experiment (three nose pokes on the same side within a 2-s inter-poke interval). Training continued until all mice performed reliably (≈1 week).

##### Sucrose preference

Two corners were assigned to 1% sucrose (both sides), and the remaining two corners provided water. Access followed the trained FR3 schedule with 90% reward probability. Preference testing ran for 2 weeks with minimal disturbance (except weekly cage changes).

##### Sucrose punishment

During punishment, sucrose access was paired with a brief air puff (1 s) delivered with 30% probability, 1.5 s after door opening. Punishment was run in two 3-day blocks, separated by 3–5 days of unpunished sucrose preference testing. *Data processing:* IntelliCage archive files were imported into R, cleaned and summarized using standard packages (archive v1.1.12, xml2 v1.3.8, tidyverse v2.0.0). Summary data were plotted in GraphPad Prism 9/10.

### Statistical analysis

Statistical analyses were performed in Prism (GraphPad). Unless otherwise stated, two-group comparisons used unpaired two-tailed t-tests. Experiments with repeated measures were analyzed using two-way repeated-measures ANOVA, and other designs used one-way ANOVA or paired t-tests, as appropriate. For post hoc analysis, we used Dunnett’s multiple-comparison test, comparing all experimental groups against the control group only, unless otherwise stated. Full details of statistical tests, degrees of freedom, exact p values and sample sizes are provided in Table S1. Data are presented as mean ± s.e.m., and individual animal data are shown whenever feasible. Statistical significance was defined as α = 0.05; * denotes p < 0.05, # denotes a trend (0.05 < p < 0.1), and ns denotes p ≥ 0.05. Sample sizes were guided by prior studies using comparable approaches and were chosen to detect robust effects while minimizing animal use.

## ACKNOWLEDGMENT

This research was supported by the Intramural Research Program of the National Institutes of Health to VAA (ZIA AA000421, ZIA MH002987), to MM (ZIA DA000069). We would like to thank the NIMH IRP Rodent Behavioral Core for their help with behavioral testing and Daniel da Silva for data analysis advice. The contributions of the NIH authors were made as part of their official duties as NIH federal employees, are in compliance with agency policy requirements, and are considered Works of the United States Government. However, the findings and conclusions presented in this paper are those of the authors and do not necessarily reflect the views of the NIH or the U.S. Department of Health and Human Services.

## AUTHOR CONTRIBUTION

Conceptualization: VAA and EMM. Experimentation: autoradiography: EV, AT, EMM, MEB and MM; dopamine voltammetry: HJS and SAL; behaviors and traits: EMM, MEB, LGA and SC; cocaine IVSA: EMM, DUD and RB. Data analysis: EMM, RB, DUD, EV, AT, LGA, SC, HJS, SAL, MM and VAA. Writing - Original Draft: EMM, VAA and RB. Review & Editing: all authors; Supervision: VAA and MM. Funding Acquisition: VAA and MM.

## COMPETING INTEREST DECLARATION

The authors declare no competing interests.

## DATA AND CODE AVAILABILTY

Data is available in a public repository Mendeley DOI: 10.17632/3zwyv4grzz.1.

**Supplementary Information is available for this paper.**

## Notes

### Competing Interest Statement

The authors have declared no competing interest.

