## Extended Data Figures for "D2 autoreceptors gate vulnerability to cocaine use disorder"

### Supplementary Figure 1 - Murray et al.

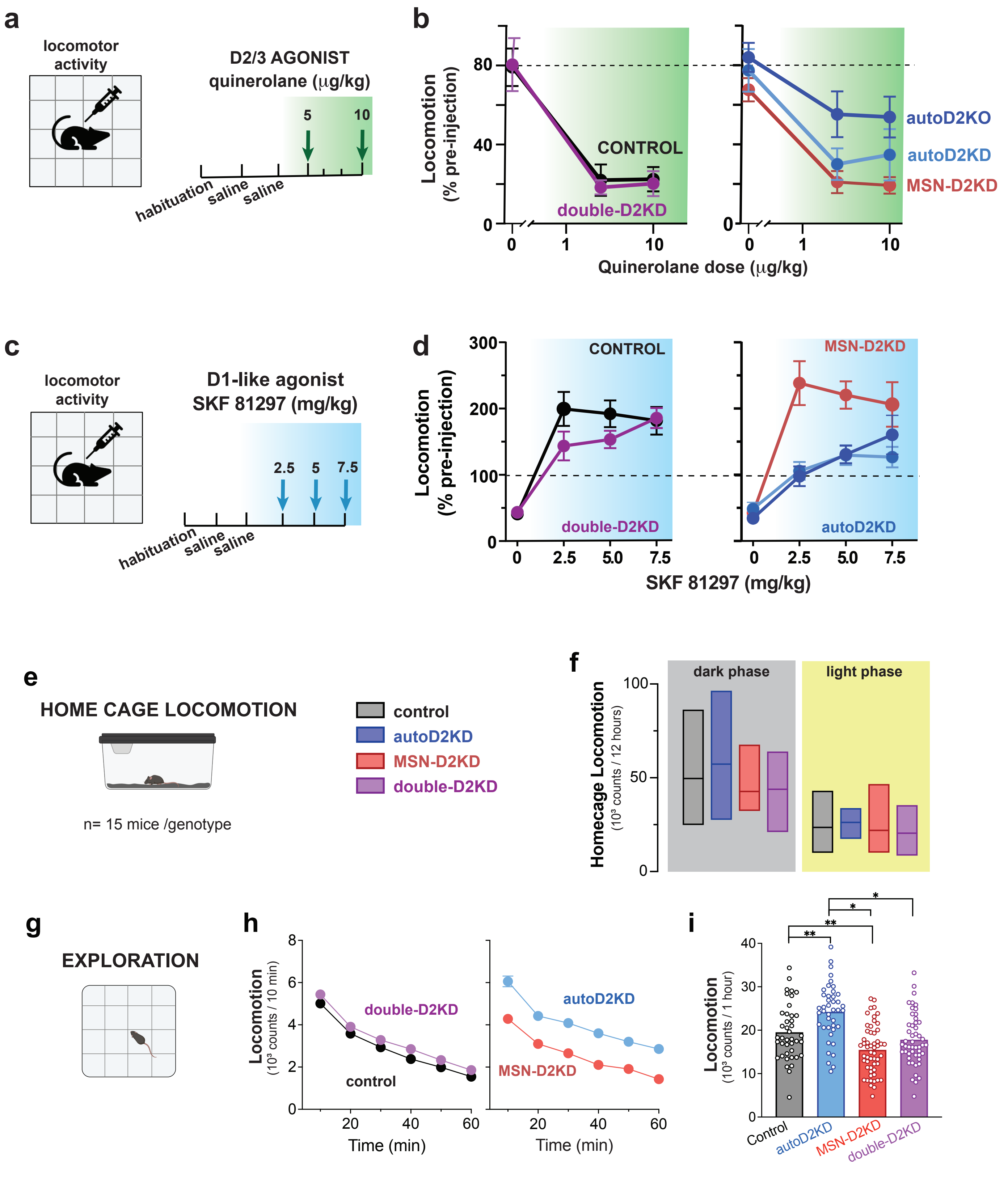

**Supplementary Fig. 1. Basal locomotion and responses to dopamine agonists.**

**a,c,** Experimental timelines for assessing dose–response effects of the D2/3 agonist quinelorane (5, 10 µg/kg, i.p.) (**a**) and the D1-like agonist SKF81297 (2.5, 5, 7.5 mg/kg, i.p.) (**c**) in locomotor activity chambers. Most mice were tested with both agonists, with a 2-week washout; SKF81297 was tested first.

**b**, **d**, Locomotor activity following saline or agonist injection, expressed as percent of each mouse's pre-injection baseline for quinelorane (**b**) * main effect of dose: $F(1.83,43.81) = 54.22$, $p < 0.0001$; no genotype $F(3,67) = 0.8$ $p = 0.49$; and for SKF81297(**d**), main effect of dose: $F(2.373, 178.8) = 46.98$, $p <0.0001$; main effect of genotype: $F(4, 80) = 6.366$, $p = 0.0002$). Left plot for controls (black) and double-D2KD (purple). Right plot for autoD2KD (blue), auto-D2KO (dark blue), and MSN-D2KD (red). n = 7–15 mice per genotype per dose. Symbols indicate group means; error bars, ± s.e.m. *$p \leq 0.05$ versus control.

**e,** Schematic of the single-housed home-cage locomotor monitoring setup.

**f,** Basal locomotion during the light and dark phases (n = 15 mice per genotype). Box plots show min–max; center line indicates the mean.

**g,** Novelty-driven exploration assessed in a novel open-field arena.

**h,** Time course of novelty-induced locomotion (10-min bins) for littermate mice of each genotype (n = 41 - 53).

**i**, Total novelty-induced locomotion over 60 min. Points show individual mice; bars indicate mean ± s.e.m. **$p \leq 0.01$ versus control.

#### Supplementary Figure 2 - Murray et al.

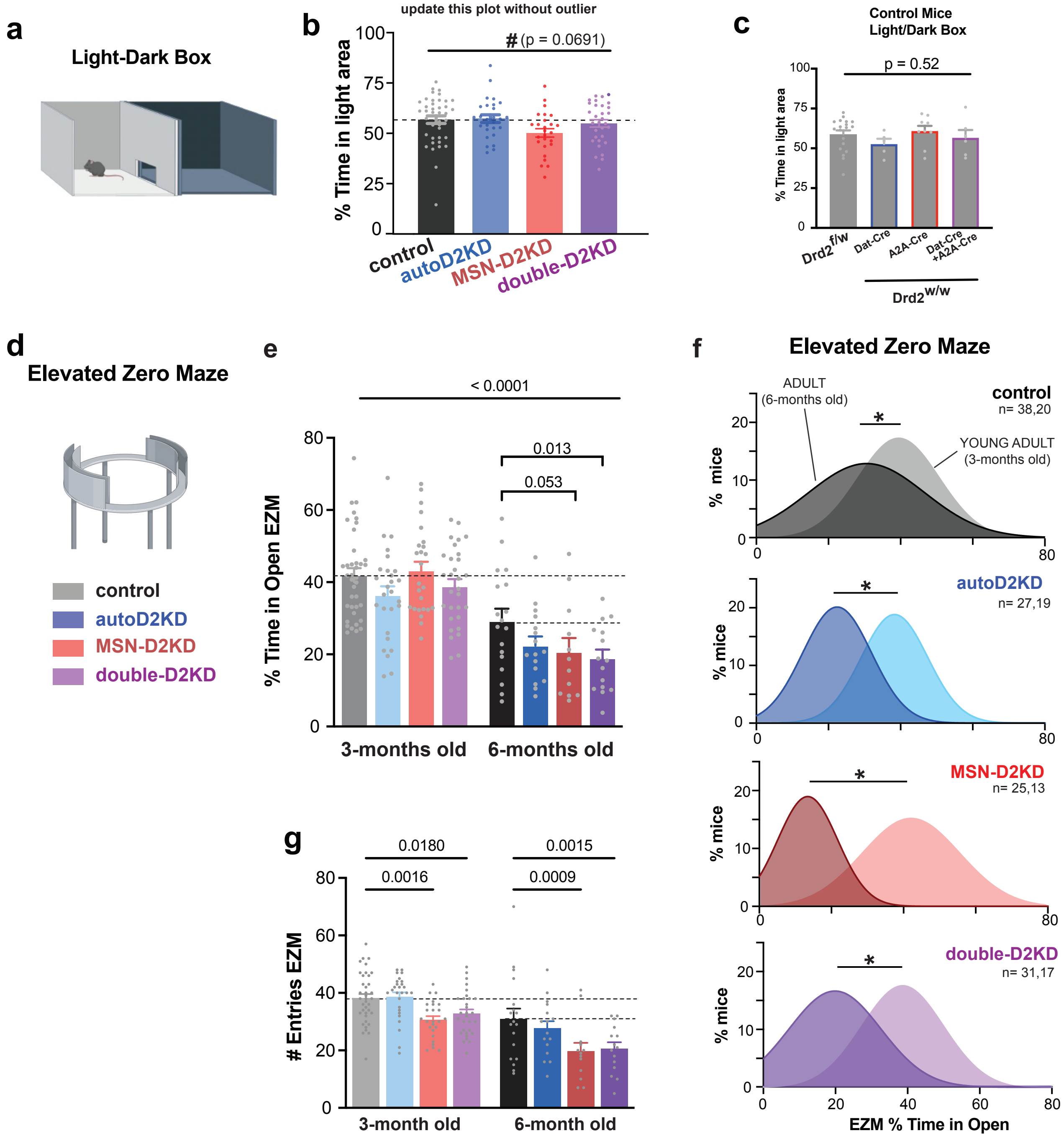

**Supplementary Fig. 2. Risk-avoidance behavior.**

**a,** Schematic of the light–dark box.

**b,** Percent time spent in the illuminated compartment for littermate mice of each genotype (n = 38 control, 27 autoD2KD, 25 MSN-D2KD and 31 double-D2KD). Points indicate individual mice; bars show mean ± s.e.m. #, trend for a genotype effect (one-way ANOVA, p = 0.069).

**c,** Light–dark box performance in Cre-driver control lines $Drd2^{f/w}$, $Drd2^{w/w}$; $DAT^{IRES\text{-}Cre}$, $Drd2^{w/w}$;Adora2A-$Cre^{+/-}$, $Drd2^{w/w}$;$DAT^{IRES\text{-}Cre}$; Adora2a-$Cre^{+/-}$ show no differences between control types (one-way ANOVA, F (3, 35) = 0.769, p = 0.518).

**d,** Schematic of the elevated zero maze (EZM).

**e,g,** EZM performance at 3 and 6 months of age: percent time in open quadrants (**e**) and number of open-quadrant entries (**g**) for each genotype (3 months / 6 months: controls n = 38 / 20; autoD2KD n = 27 / 19; MSN-D2KD n = 25 / 13; double-D2KD n = 31 / 17). Points indicate individual mice; bars show mean ± s.e.m. Two-way ANOVA main effect of age, p = 0.0001; post hoc p values are indicated.

**f,** Frequency distributions of time spent in the illuminated compartment in the light–dark box at 3 months (light shading) and 6 months (dark shading) of age. *p ≤ 0.05.

#### Supplementary Figure 3 - Murray et al.

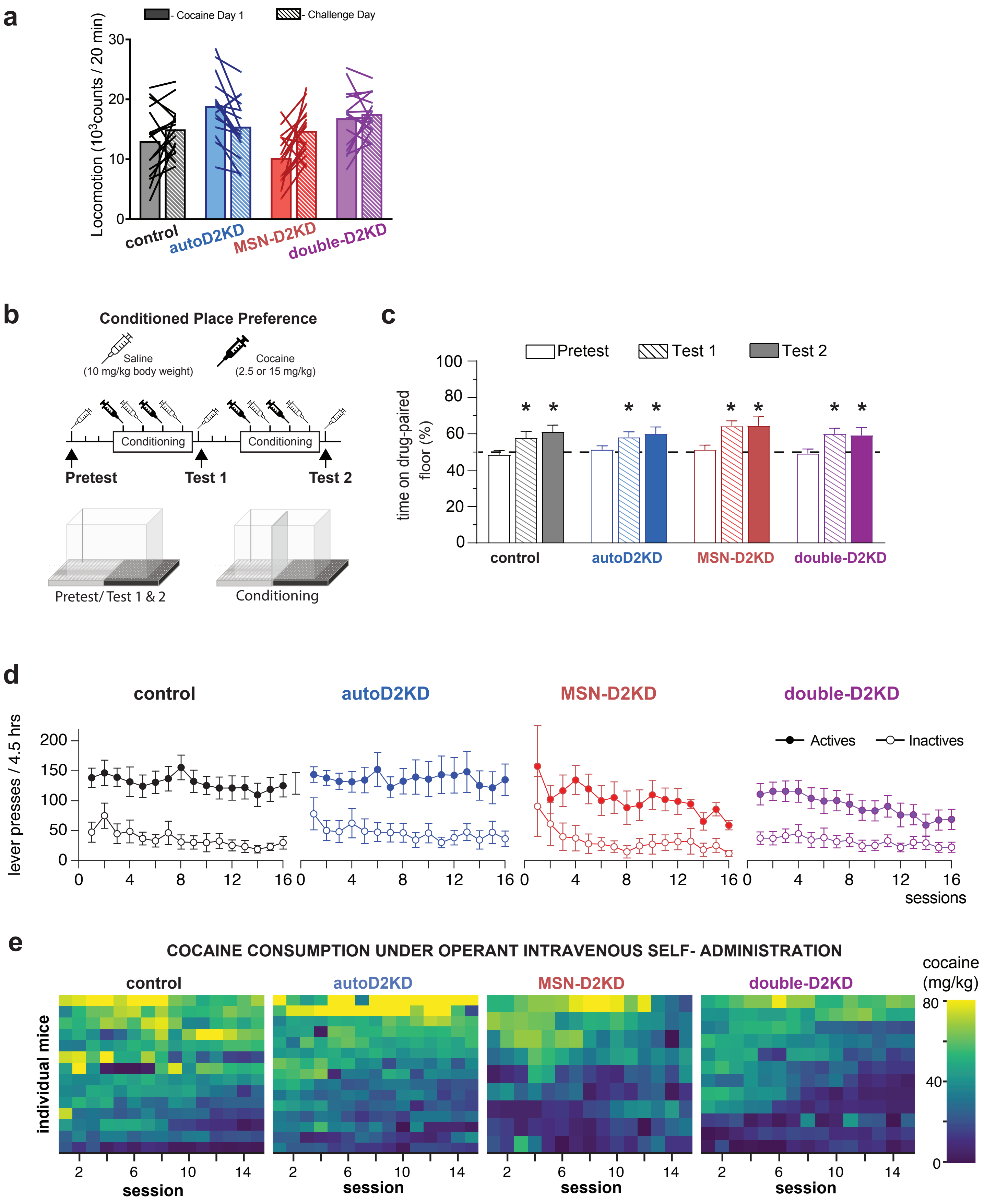

**Supplementary Fig. 3. Acute and repeated cocaine responses.**
**a,** Cocaine-induced locomotion (20-min epoch) during the first cocaine exposure (day 1) and during the cocaine challenge after 2 weeks of abstinence (controls, n = 16; autoD2KD, n = 17; MSN-D2KD, n = 15; double-D2KD, n = 15). Points show individual mice; lines connect paired measurements within mice; bars indicate mean ± s.e.m.
**b,** Experimental timeline and schematic of the conditioned place preference (CPP) apparatus.
**c,** Percentage of time spent on the cocaine-paired floor during the pre-test (open bars), Test 1 (striped bars) and Test 2 (filled bars) of conditioned place preference (n = 13-15 mice per genotype). Bars indicate mean ± s.e.m.; $^*p \leq 0.05$.
**d,** Lever-press rates on the active (filled symbols) and inactive (open symbols) levers across sessions for each genotype (controls, n = 13; autoD2KD, n = 15; MSN-D2KD, n = 9; double-D2KD, n = 12). Symbols indicate mean ± s.e.m.
**e,** Heat maps of daily cocaine intake (mg $kg^{-1}$) across 16 IVSA sessions for individual mice in each genotype; color scale denotes intake.

#### Supplementary Figure 4 - Murray et al.

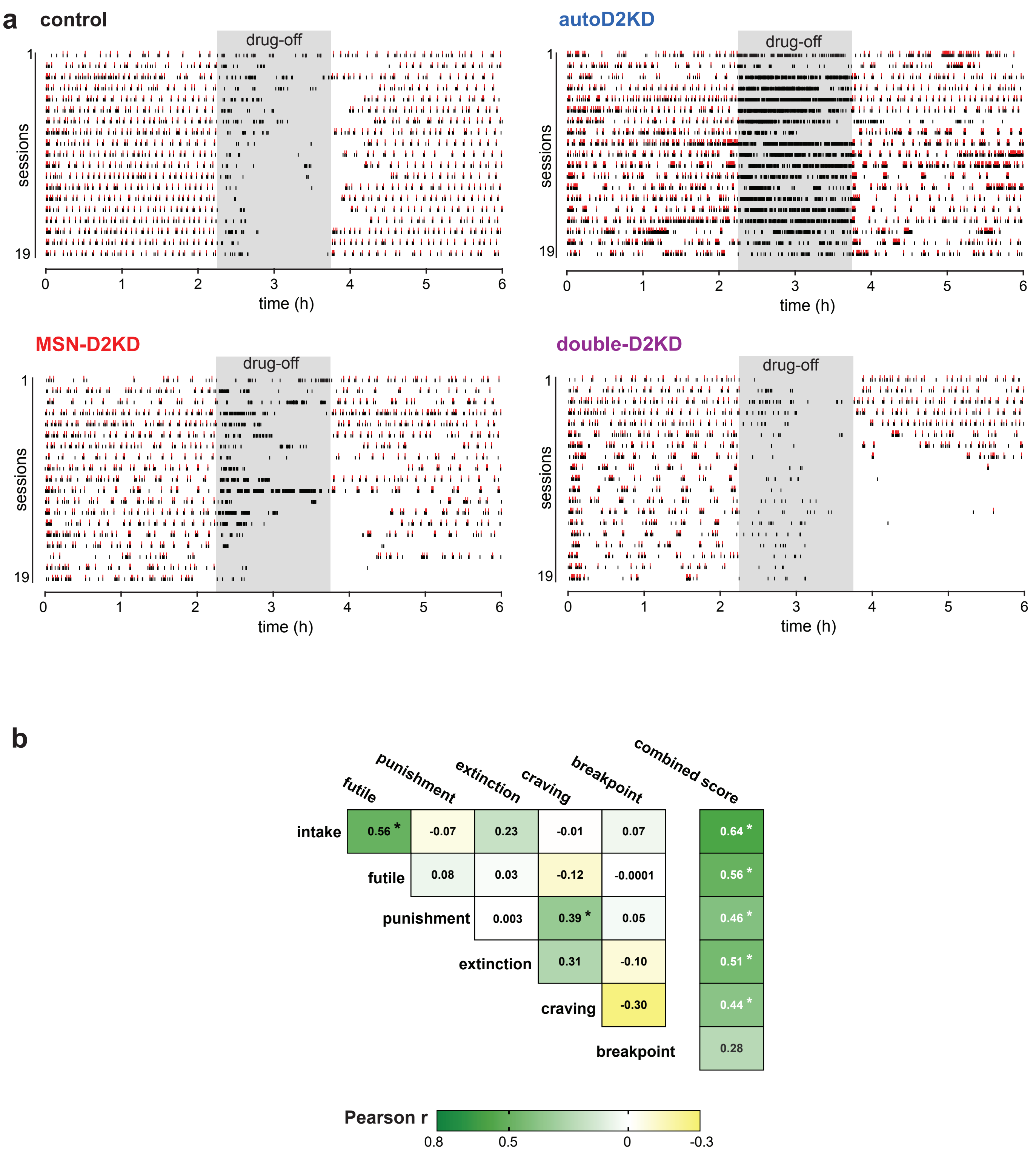

**Supplementary Fig. 4. Cocaine intravenous self-administration enables multi-domain assessment of addictive-like behaviors.**

**a**, Representative rater plots of cocaine IVSA during 19 sessions after acquisition. Black ticks indicate active-lever presses and red ticks indicate earned cocaine infusions (1 mg/kg/infusion) under FR3 schedule. Each 6-hour session included a 90 min signaled ("drug-off") period (shaded), during which responses had no programmed consequences and were quantified as futile responding.

**b**, Correlation matrix across cocaine-related behavioral measures. Colors indicate Pearson's r; * $p < 0.05$.

### Supplementary Figure 5 - Murray et al.

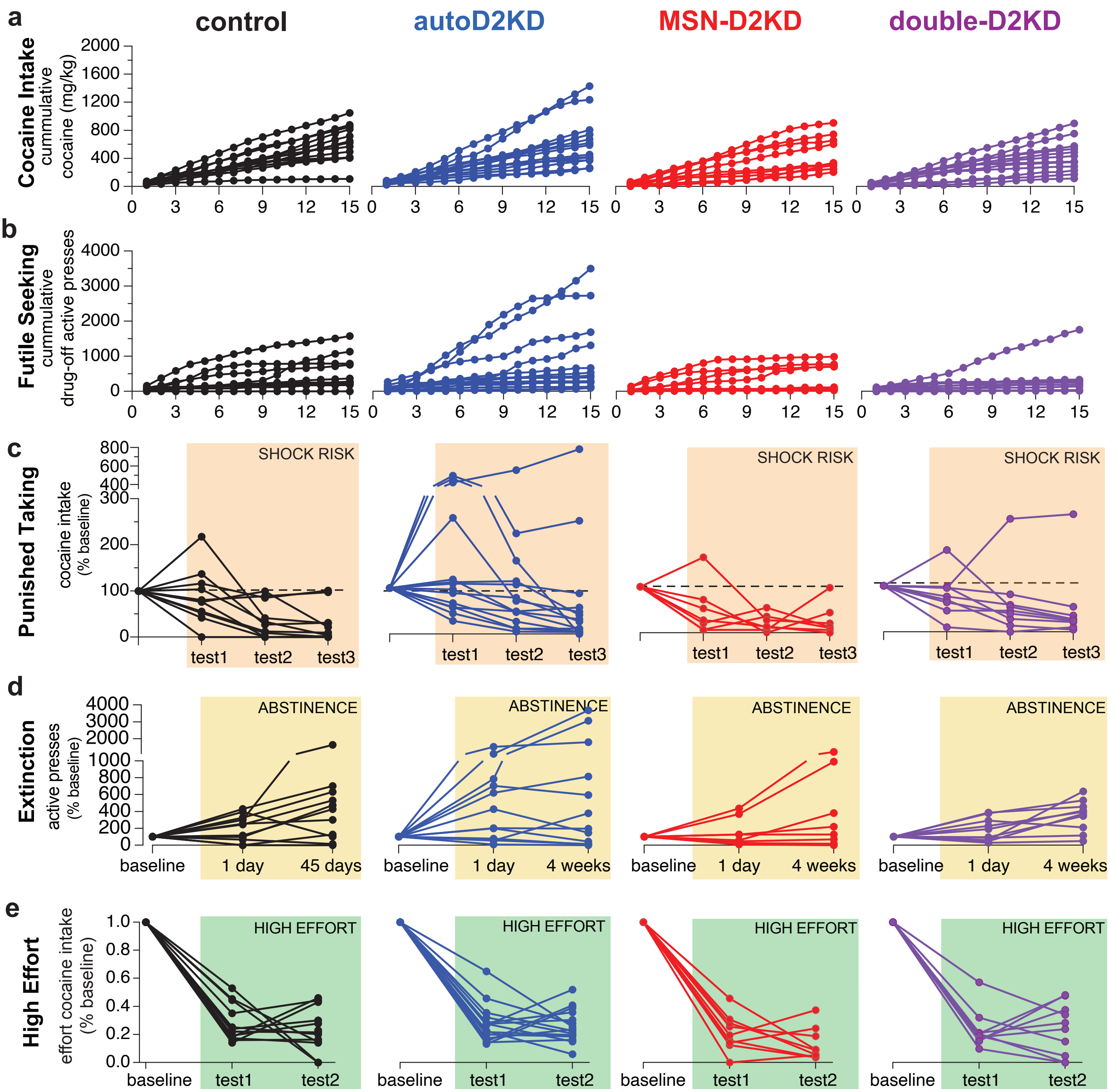

**Supplementary Fig. 5. autoD2KD mice show increased inter-individual variability in cocaine-related behaviors during IVSA.**

**a,b,** Across-session trajectories for cumulative cocaine intake (a) and futile responding during the signaled drug-unavailable (“drug-off”) period (b) over 16 IVSA sessions for controls (n = 13, black), autoD2KD (n = 15, blue), MSN-D2KD (n = 9, red) and double-D2KD (n = 12, purple). Symbols indicate individual mice.

**c–e,** Individual performance during foot-shock punishment across three tests (c), cue/context-driven seeking during abstinence (d), and high-effort responding assessed with progressive-ratio tests (e). Data are normalized to each mouse’s baseline (pre-punishment intake, pre-abstinence responding, or FR3 performance, respectively).

#### Supplementary Figure 6- Murray et al.

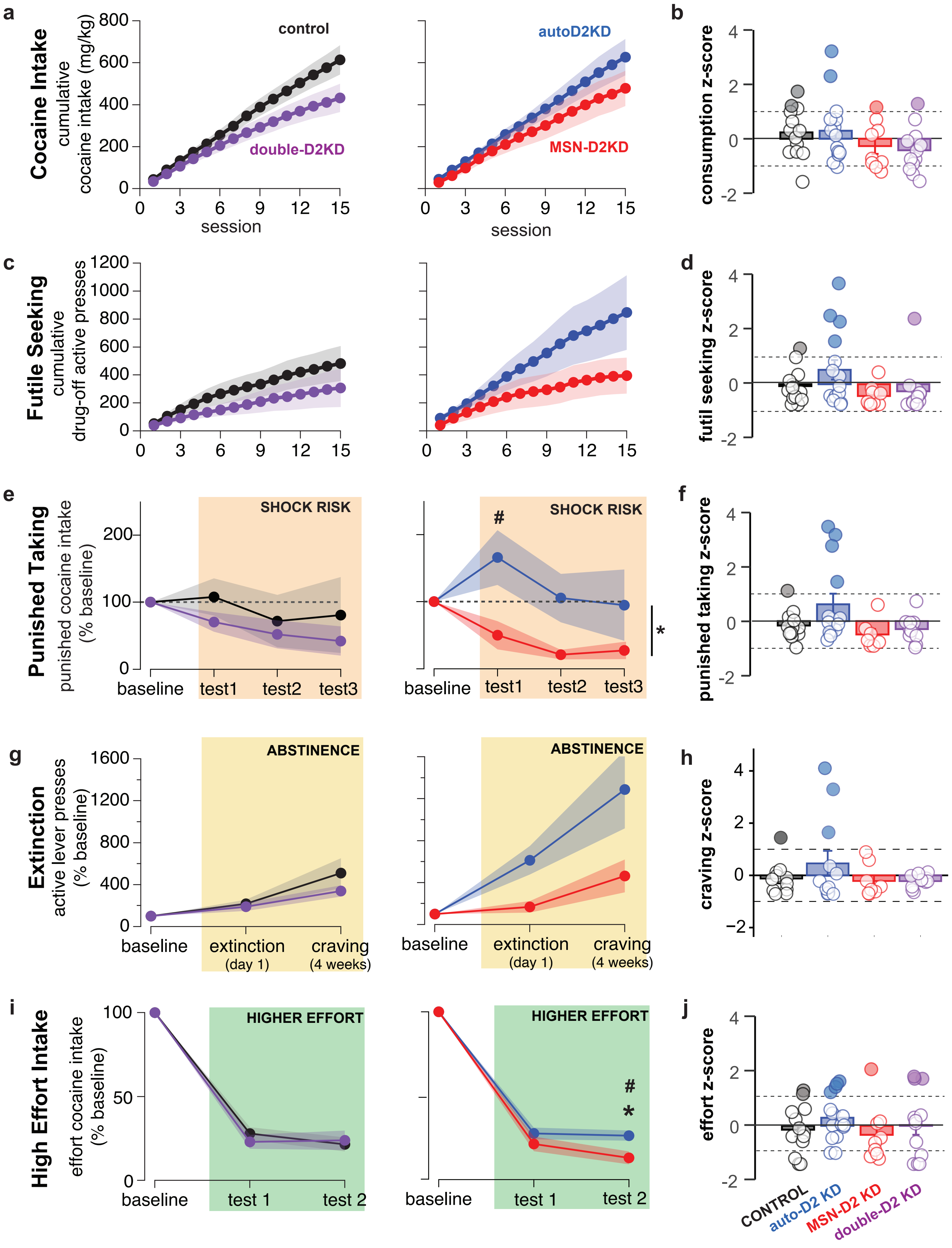

**Supplementary Fig. 6. Cocaine IVSA behavioral measures and z-score across genotypes.**
**a,c,e,g,i,** Mean behavioral measures by genotype for cumulative cocaine intake (**a**), cumulative futile responding (**c**), punished cocaine consumption (**e**), extinction and craving responding (**g**), and high-effort consumption (**i**). For each measure, the left panel shows controls (black) and double-D2KD (purple), and the right panel shows autoD2KD (blue) and MSN-D2KD (red). Symbols indicate group means; shaded bands show ± s.e.m.

**b,d,f,h,j,** Z-scores by genotype for cocaine consumption (**b**), futile seeking (**d**), punished cocaine consumption (**f**), craving (**h**), and high-effort intake (**j**). Points indicate individual mice; bars show mean.

For all panels, *$P \leq 0.05$ and #$0.05 < P \leq 0.1$ versus control.

#### Supplementary Figure 7- Murray et al.

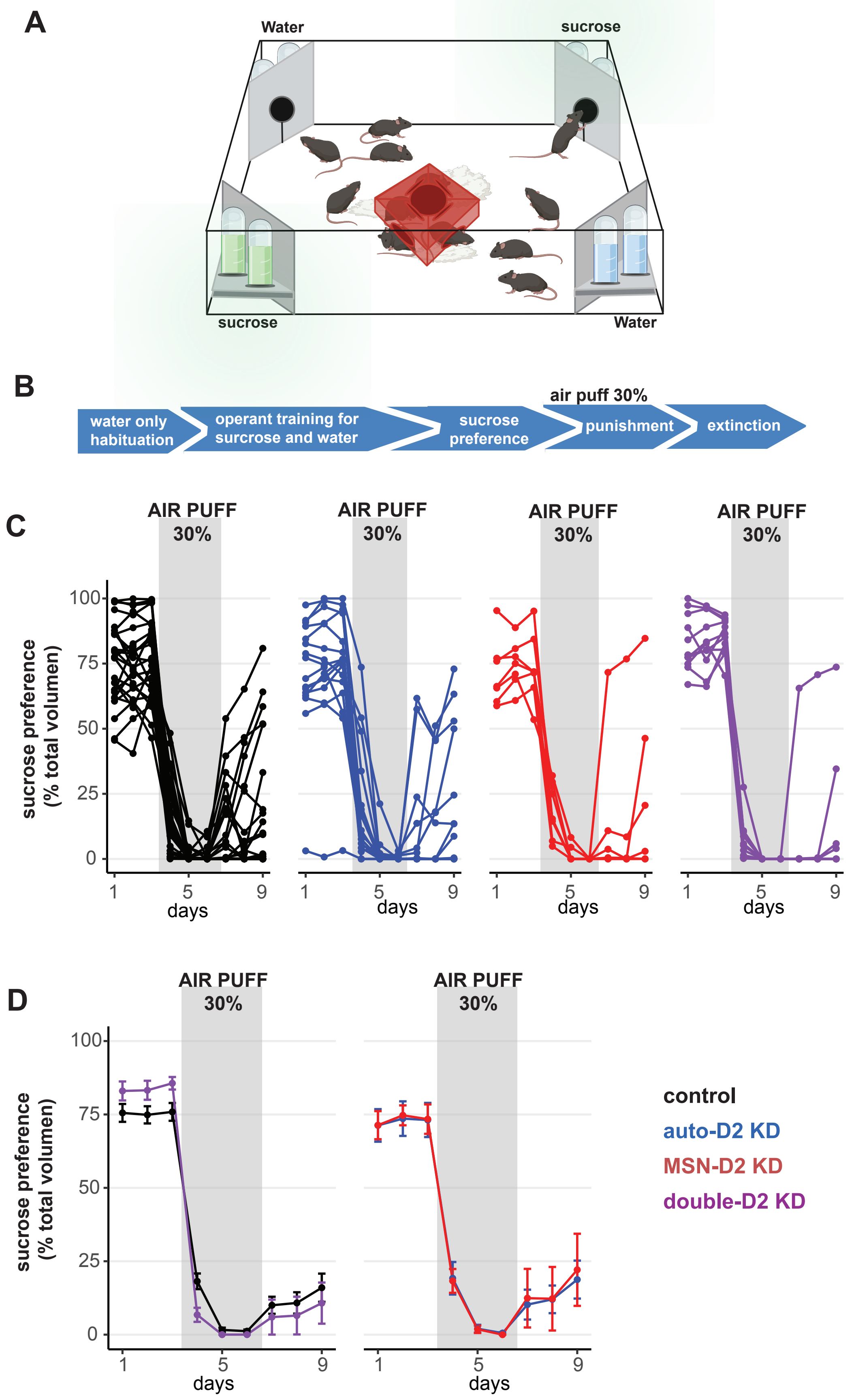

**Supplementary Fig. 7. Intact sucrose preference and punishment sensitivity across genotypes**

**a,** Schematic of the IntelliCage system for self-paced operant testing in socially housed mice. **b,** Experimental timeline showing operant access to water and 1% sucrose, followed by a 3-day punishment phase in which 30% of sucrose-access events were paired with an air puff (2 s, psi). **c,d,** Sucrose preference (sucrose licks / water licks) shown for individual mice (**c**) and as genotype means (**d**) for controls (black), autoD2KD (blue), MSN-D2KD (red) and double-D2KD (purple).
