## Supplementary material for "D2 autoreceptors gate vulnerability to cocaine use disorder": STATISTIC TABLE

Murray et al. Statistical Anlaysis Data Table File

| Figure | Measurement | # of mice | Statistical test | Results | p value |
| --- | --- | --- | --- | --- | --- |
| 1b | Ddr2 mRNA expression | n = 3 - 5 | **Region** | | |
| | | | Interaction (Brain Region x Genotype) | F (6, 28) = 1.249 | 0.3121 |
| | | | Region | F (1.247, 17.46) = 5.341 | 0.027 |
| | | | Genotype | F (3, 16) = 12.81 | 0.0002 |
| | | | **Dunnett's multiple comparisons test (region)** | | |
| | | | *Cortex* | | |
| | | | Control vs. double-D2KD | | 0.7593 |
| | | | Control vs. Auto-D2KD | | 0.9496 |
| | | | Control vs. MSN-D2KD | | 0.6129 |
| | | | *NAc* | | |
| | | | Control vs. double-D2KD | | 0.021 |
| | | | Control vs. Auto-D2KD | | 0.418 |
| | | | Control vs. MSN-D2KD | | 0.0192 |
| | | | *Dorsal Striatum* | | |
| | | | Control vs. double-D2KD | | 0.001 |
| | | | Control vs. Auto-D2KD | | 0.3019 |
| | | | Control vs. MSN-D2KD | | 0.0002 |
| 1d | D2-like receptor radioligand binding | n = 3 - 9 | **Region** | | |
| | | | Interaction (Brain Region x Genotype) | F (3, 32) = 5.865 | 0.0026 |
| | | | Region | F (1, 32) = 11.11 | 0.0022 |
| | | | Genotype | F (3, 32) = 43.36 | <0.0001 |
| | | | **Dunnett's multiple comparisons test (region)** | | |
| | | | *Dorsal Striatum* | | |
| | | | Control vs. double-D2KD | | <0.0001 |
| | | | Control vs. Auto-D2KD | | 0.0041 |
| | | | Control vs. MSN-D2KD | | <0.0001 |
| | | | *NAc* | | |
| | | | Control vs. double-D2KD | | 0.0037 |
| | | | Control vs. Auto-D2KD | | 0.064 |
| | | | Control vs. MSN-D2KD | | <0.0001 |
| | | | **2W ANOVA - Genotype x Region** | | |
| | | | Interaction (Genotype x Region) | F (3, 12) = 5.122 | 0.0165 |

| | | | | | |
|---|---|---|---|---|---|
| | | | Genotype | F (3, 12) = 5.995 | 0.0098 |
| | | | Region | F (1, 12) = 10.74 | 0.0066 |
| | | | **Tukey's multiple comparisons test (region)** | | |
| | | | *Dorsal Striatum* | | |
| | | | double-D2KD vs. AutoD2 | | 0.984 |
| | | | AutoD2 vs. Control | | 0.8211 |
| | | | AutoD2 vs. MSN | | 0.1082 |
| 1f | D1-like receptor radioligand binding | n = 3 - 6 | double-D2KD vs. Control | | 0.5839 |
| | | | double-D2KD vs. MSN | | 0.0496 |
| | | | MSN vs. Control | | 0.269 |
| | | | *NAc* | | |
| | | | double-D2KD vs. AutoD2 | | 0.7971 |
| | | | AutoD2 vs. Control | | 0.0183 |
| | | | AutoD2 vs. MSN | | 0.0006 |
| | | | double-D2KD vs. Control | | 0.1653 |
| | | | double-D2KD vs. MSN | | 0.007 |
| | | | MSN vs. Control | | 0.2445 |

| sig? |
| --- |
| * |
| *** |
| ns |
| ns |
| ns |
| * |
| ns |
| * |
| *** |
| ns |
| *** |
| ** |
| ** |
| *** |
| **** |
| ** |
| **** |
| ** |
| # |
| **** |
| * |

**
**

ns
ns
ns
ns
*
ns

ns
*
***
ns
**
ns

| Figure | Measurement | # of mice | Statistical test |
|---|---|---|---|
| 2b | D1/D2 Receptor Ratio (Radioligand binding) | n = 3 - 6 | **2W mixed-effects model - Genotype x Region**<br>Interaction (Region x Genotype)<br>Region<br>Genotype<br>**Dunnett's multiple comparisons test**<br>*Dorsal Striatum*<br>Control vs. double-D2KD<br>Control vs. Auto-D2KD<br>Control vs. MSN-D2KD<br>*NAc*<br>Control vs. double-D2KD<br>Control vs. Auto-D2KD<br>Control vs. MSN-D2KD |
| 2d | Locomotor Response to 5 µg/kg Quinelorane (D2-like agonist) | n = 6 - 11 | **One-way ANOVA** |
| 2d | D1-like (5 mg/kg) and D2/3-like (5 ug/mg) agonist locomotor response | n = 6 - 21 | **2W ANOVA - Genotype x Agonist Type**<br>Interaction (Genotype x Agonist Type)<br>Genotype<br>Agonist Type<br>**Dunnett's multiple comparisons test**<br>*D2/3-like agonist (quinelorane)*<br>Control vs. Auto-D2KD<br>Control vs. MSN-D2KD<br>Control vs. double-D2KD<br>*D1-like agonist (SKF81297)*<br>Control vs. Auto-D2KD<br>Control vs. MSN-D2KD<br>Control vs. double-D2KD |

| Results | p value | Significance? |
|---|---|---|
| F (3, 25) = 7.258 | 0.0012 | ** |
| F (1, 25) = 0.2645 | 0.6115 | ns |
| F (3, 25) = 22.74 | <0.0001 | **** |
| | <0.0001 | **** |
| | 0.4828 | ns |
| | <0.0001 | **** |
| | 0.8612 | ns |
| | 0.7579 | ns |
| | 0.0002 | *** |
| F (3, 31) = 0.6602 | 0.5828 | ns |
| F (3, 97) = 2.894 | 0.0392 | * |
| F (3, 97) = 2.020 | 0.1162 | ns |
| F (1, 97) = 56.21 | <0.0001 | **** |
| | 0.9869 | ns |
| | 0.9997 | ns |
| | >0.9999 | ns |
| | 0.0165 | * |
| | 0.4409 | ns |
| | 0.1879 | ns |

| Figure | Measurement | # of mice | Statistical test | Results | p value | Significance? |
|---|---|---|---|---|---|---|

| Figure | Measurement |
|---|---|
| 4c, S1h-i | Open Field Locomotion: Novel Exploration |
| 4d, S2b | Light-Dark Box: Duration in Light Area |

| # of mice | Statistical test |
| --- | --- |
| | **One-way ANOVA** |
| | **Tukey's multiple comparisons test** |
| n = 41 - 53 | DbHET vs. AutoD2 |
| | DbHET vs. MSN-D2KD |
| | DbHET vs. Control |
| | AutoD2 vs. MSN-D2KD |
| | AutoD2 vs. Control |
| | MSN-D2KD vs. Control |
| | **One-way ANOVA** |
| | **Dunnett's multiple comparisons test** |
| n = 25 - 38 | Control vs. double-D2KD |
| | Control vs. Auto-D2KD |
| | Control vs. MSN-D2KD |

| Results | p value | Significance? |
|---|---|---|
| F (3, 183) = 17.28 | <0.0001 | **** |
| | <0.0001 | **** |
| | 0.2077 | ns |
| | 0.5304 | ns |
| | <0.0001 | **** |
| | 0.0029 | ** |
| | 0.0079 | ** |
| F (3, 119) = 3.32 | | |
| | 0.5116 | ns |
| | 0.9924 | ns |
| | 0.0107 | * |

| Figure | Measurement | # of mice |
|---|---|---|
| 5b | Cocaine Sensitization: Habitutation Locomotor Response | n = 16 - 18 |
| 5b | Cocaine Sensitization: comparisons across saline day 1 through cocaine day 5 | n = 16 - 18 |

| | | |
|---|---|---|
| 5c | <u>Cocaine Sensitization: Comparison of Cocaine Sensitization Score</u> | n = 16 - 18 |

| Statistical test | Results | p value | ? |
|---|---|---|---|
| **One-way ANOVA** | F (3, 63) = 5.281 | 0.0026 | ** |
| **Dunnett's multiple comparisons test** | | | |
| Control vs. Auto-D2KD | | 0.2863 | ns |
| Control vs. MSN-D2KD | | 0.0746 | ns |
| Control vs. double-D2KD | | 0.5735 | ns |
| **2W mixed-effects model - Session Day x Genotype** | | | |
| Interaction (Cocaine x Genotype) | F (15.32, 317.5) = 3.874 | <0.0001 | **** |
| Session Day | F (5.107, 317.5) = 117.6 | <0.0001 | **** |
| Genotype | F (3, 63) = 6.183 | 0.0009 | *** |
| **Dunnett's multiple comparisons test** | | | |
| *Saline Day* | | | |
| Control vs. double-D2KD | | 0.8983 | ns |
| Control vs. Auto-D2KD | | 0.0508 | ns |
| Control vs. MSN-D2KD | | 0.8562 | ns |
| *Cocaine Day 1* | | | |
| Control vs. double-D2KD | | 0.1286 | ns |
| Control vs. Auto-D2KD | | 0.0123 | * |
| Control vs. MSN-D2KD | | 0.4152 | ns |
| *Cocaine Day 2* | | | |
| Control vs. double-D2KD | | 0.0913 | ns |
| Control vs. Auto-D2KD | | 0.0076 | ** |
| Control vs. MSN-D2KD | | 0.8944 | ns |
| *Cocaine Day 3* | | | |
| Control vs. double-D2KD | | 0.0243 | * |
| Control vs. Auto-D2KD | | 0.7034 | ns |
| Control vs. MSN-D2KD | | 0.8636 | ns |
| *Cocaine Day 4* | | | |
| Control vs. double-D2KD | | 0.0092 | ** |
| Control vs. Auto-D2KD | | 0.4477 | ns |
| Control vs. MSN-D2KD | | 0.9963 | ns |
| *Cocaine Day 5* | | | |
| Control vs. double-D2KD | | 0.0355 | * |

| | | | |
|---|---|---|---|
| Control vs. Auto-D2KD | | 0.4497 | ns |
| Control vs. MSN-D2KD | | 0.7022 | ns |
| *Challenge Day* | | | |
| Control vs. double-D2KD | | 0.0768 | ns |
| Control vs. Auto-D2KD | | 0.9813 | ns |
| Control vs. MSN-D2KD | | 0.861 | ns |
| **One-way ANOVA** | F (3, 61) = 4.45 | 0.0068 | ** |
| **Tukey's multiple comparisons test** | | | |
| double-D2KD vs. Auto-D2KD | | 0.3533 | ns |
| double-D2KD vs. MSN-D2KD | | 0.4339 | ns |
| Control vs. double-D2KD | | 0.4633 | ns |
| Auto vs. MSN-D2KD | | 0.0141 | * |
| Control vs. Auto-D2KD | | 0.015 | * |
| Control vs. MSN-D2KD | | 0.9999 | ns |

| Figure | Measurement | # of mice | Statistical test |
|---|---|---|---|
| 6c | IVSA: genotype rate of aquisition | n = 9 - 15 | **Chi-Square** |
| 6d | IVSA: days to task aquisition | n = 9 - 15 | **One-way ANOVA** |
| 6f | IVSA: cummulative cocaine intake | | **Two-way RM ANOVA** |
| | | | sessions x Genotype |
| | | n = 13 - 22 | sessions |
| | | | Genotype |
| 6g | IVSA: total cocaine consumption | n = 13 - 22 | **Unpaired T-test** |
| 6h | IVSA: cummulative futile responses | n = 13 - 22 | **Two-way RM ANOVA** |
| | | | sessions x Genotype |
| | | | sessions |
| | | | Genotype |
| | | | Subject |
| 6i | IVSA: total futile responses | n = 13 - 22 | **One-way ANOVA** |
| | | | autoD2KD vs MSN-D2KDcombined |
| 6j | IVSA: punished consumption | n = 13 - 22 | **Two-way RM ANOVA** |
| | | | Punishment |
| | | | Genotype |
| | | | Punishment x Genotype |
| 6k | IVSA: first punishment test | n = 13 - 22 | **One-way ANOVA** |
| | | | autoD2KD vs MSN-D2KDcombined |
| 6l | IVSA: extinction and craving | n = 13 - 22 | **Two-way RM ANOVA** |
| | | | Abstinence x Genotype |
| | | | Abstinence |
| | | | Genotype |
| | | | Subject |
| 6m | IVSA: extinction | n = 13 - 22 | **One-way ANOVA** |
| | | | autoD2KD vs MSN-D2KDcombined |
| | | | control vs autoD2KD |
| 6n | IVSA: high effort consumption | n = 13 - 22 | **Two-way RM ANOVA** |
| | | | High Effort x Genotype |
| | | | High Effort |
| | | | Genotype |
| | | | Subject |
| 6o | IVSA: z-scores | n = 13 - 22 | **Mixed-effects model (REML)** |
| | | | Genotypes |
| | | | Behaviors |
| | | | Genotypes x Behaviors |

| Results | p value | ? |
|---|---|---|
| X² = 0.24 | 0.97 | ns |
| (3, 45) = 0.215 | 0.8852 | ns |
| F (28, 644) = 1.[illegible] | P=0.0036 | Yes |
| F (1.120, 51.53[illegible] | P<0.0001 | Yes |
| F (2, 46) = 1.91 | P=0.1595 | ns |
| t (34) = 1.8 | 0.075 | # |
| F (28, 672) = 2.[illegible] | P=0.0012 | yes |
| F (1.149, 55.14[illegible] | P<0.0001 | yes |
| F (2, 48) = 2.81[illegible] | P=0.0700 | # |
| F (48, 672) = 3[illegible] | P<0.0001 | yes |
| F (2, 48) = 2.65[illegible] | P=0.0803 | # |
| post hoc | P=0.0741 | # |
| F (1.777, 74.64[illegible] | 0.0365 | yes |
| F (2, 43) = 2.84[illegible] | 0.0693 | # |
| F (4, 84) = 2.16[illegible] | 0.0797 | # |
| F (2, 42) = 3.77[illegible] | P=0.0311 | yes |
| post hoc | P=0.0234 | yes |
| F (4, 76) = 1.65 | P=0.1702 | ns |
| F (1.155, 43.87[illegible] | P=0.0004 | yes |
| F (2, 38) = 2.83[illegible] | P=0.0715 | # |
| F (38, 76) = 1.9[illegible] | P=0.0064 | yes |
| F (2, 38) = 4.68[illegible] | P=0.0151 | yes |
| post hoc | P=0.0153 | yes |
| post hoc | P=0.0656 | # |
| F (4, 86) = 0.62[illegible] | P=0.6447 | ns |
| F (2, 86) = 650.[illegible] | P<0.0001 | yes |
| F (2, 43) = 2.06 | P=0.1397 | ns |
| F (43, 86) = 0.8[illegible] | P=0.6620 | ns |
| F (2, 46) = 7.38[illegible] | 0.0017 | yes |
| F (3.620, 151.3[illegible] | 0.8154 | ns |
| F (10, 209) = 1.[illegible] | 0.4089 | ns |

| Figure | Measurement | # of mice | Statistical test | Results |
|---|---|---|---|---|
| 7a | IVSA: genotype rate of aquisition | n = 13,15,21 | **Two-way ANOVA** | |
| | | | Interaction | F (10, 264) = 0.629[illegible] |
| | | | Behaviors | F (5, 264) = 0.02231 |
| | | | Genotype | F (2, 264) = 7.624 |
| 7b | Addictive like Behaviors: z-scores | n = 13,15, 21 | **One-way ANOVA** | F (2, 46) = 6.872 |
| | | | control vs. autoD2KD | |
| | | | control vs. MSN-D2KDcombined | |
| | | | autoD2KD vs. MSN-D2KDcombined | |
| 7c | Addictive like Behaviors: Proportions | n = 13, 15, 21 | **Fisher's exact test** | |

| p value | ? |
|---|---|
| 0.7883 | ns |
| 0.9998 | ns |
| 0.0006 | yes |
| 0.0024 | yes |
| 0.0688 | # |
| 0.5315 | ns |
| 0.0017 | yes |
| 0.007 | Yes |

| Figure | Measurement | # of mice |
| --- | --- | --- |
| S1b | Controls Locomotor Dose Response to D2-like agonist (Quinelorane): 0 , 5 and 10 µg/kg | n = 6 - 15 |
| S1b | Control vs. AutoD2-KO Locomotor Dose Response to D2-like agonist (Quinelorane): 5 and 10 µg/kg | n = 6 - 16 |
| S1b | Locomotor Dose Response to D2-like agonist (Quinelorane): 0, 5 and 10 µg/kg | n = 6 - 16 |

S2d

Locomotor Dose Response to D1-like Agonist (SKF81297): 0, 2.5, 5.0, and 7.5 mg/kg

n = 15 - 21

| | | |
|---|---|---|
| SF 1f | <u>Open Field Locomotion: 24-Hour Homecage Baseline</u> | n = 45, 15 per genotype |

| Statistical test | Results | p value | Significance? |
|---|---|---|---|
| **Mixed-effects analysis** | F (1.954, 8.794) = 12.46 | 0.0028 | ** |
| **Two-way RM ANOVA** | | | |
| Interaction (Genotype x Dose) | F (1, 18) = 0.01359 | 0.9085 | ns |
| Dosage | F (1, 18) = 0.003238 | 0.9552 | ns |
| Genotype | F (1, 18) = 4.341 | 0.0517 | # |
| **Mixed-effects analysis** | | | |
| Interaction (Genotype x Dose) | F (5.476, 43.81) = 0.1983 | 0.969 | ns |
| Dosage | F (1.825, 43.81) = 54.22 | <0.0001 | **** |
| Genotype | F (3, 67) = 0.8041 | 0.496 | ns |
| **Tukey's multiple comparisons test (region)** | | | |
| 0 μg/kg (saline) vs. 0.005 μg/kg | | <0.0001 | **** |
| 0 μg/kg vs. 0.01 μg/kg | | <0.0001 | **** |
| 0.005 μg/kg vs. 0.01 μg/kg | | 0.9265 | ns |
| **2W mixed-effects model - Genotype x D1-Agonist Dose** | | | |
| Interaction (Genotype x Dose) | F (12, 226) = 2.000 | 0.0253 | * |
| Dosage | F (2.373, 178.8) = 46.98 | <0.0001 | **** |
| Genotype | F (4, 80) = 6.366 | 0.0002 | *** |
| **Dunnett's multiple comparisons test (dosage)** | | | |
| *0* | | | |
| double-D2KD vs. AutoD2 | | 0.9937 | ns |
| double-D2KD vs. MSN | | >0.9999 | ns |
| double-D2KD vs. Control | | 0.9993 | ns |
| double-D2KD vs. Auto-D2KO | | 0.8541 | ns |
| AutoD2 vs. MSN | | 0.9789 | ns |
| AutoD2 vs. Control | | 0.976 | ns |
| AutoD2 vs. Auto-D2KO | | 0.7254 | ns |
| MSN vs. Control | | 0.9999 | ns |
| MSN vs. Auto-D2KO | | 0.8148 | ns |
| Control vs. Auto-D2KO | | 0.9671 | ns |

| | | |
|---|---|---|
| *2.5* | | |
| double-D2KD vs. AutoD2 | 0.5877 | ns |
| double-D2KD vs. MSN | 0.1419 | ns |
| double-D2KD vs. Control | 0.4711 | ns |
| double-D2KD vs. Auto-D2KO | 0.4271 | ns |
| AutoD2 vs. MSN | 0.0081 | ** |
| AutoD2 vs. Control | 0.0281 | * |
| AutoD2 vs. Auto-D2KO | 0.9946 | ns |
| MSN vs. Control | 0.8834 | ns |
| MSN vs. Auto-D2KO | 0.0051 | ** |
| Control vs. Auto-D2KO | 0.0171 | * |
| *5* | | |
| double-D2KD vs. AutoD2 | 0.7529 | ns |
| double-D2KD vs. MSN | 0.0688 | # |
| double-D2KD vs. Control | 0.5101 | ns |
| double-D2KD vs. Auto-D2KO | 0.7478 | ns |
| AutoD2 vs. MSN | 0.0093 | ** |
| AutoD2 vs. Control | 0.1253 | ns |
| AutoD2 vs. Auto-D2KO | >0.9999 | ns |
| MSN vs. Control | 0.8627 | ns |
| MSN vs. Auto-D2KO | 0.0079 | ** |
| Control vs. Auto-D2KO | 0.1175 | ns |
| *7.5* | | |
| double-D2KD vs. AutoD2 | 0.0738 | # |
| double-D2KD vs. MSN | 0.9805 | ns |
| double-D2KD vs. Control | 0.9999 | ns |
| double-D2KD vs. Auto-D2KO | 0.937 | ns |
| AutoD2 vs. MSN | 0.2281 | ns |
| AutoD2 vs. Control | 0.2478 | ns |
| AutoD2 vs. Auto-D2KO | 0.8453 | ns |
| MSN vs. Control | 0.9716 | ns |
| MSN vs. Auto-D2KO | 0.8412 | ns |
| Control vs. Auto-D2KO | 0.9751 | ns |

| 2W RW ANOVA - Genotype x Phase | | | |
|---|---|---|---|
| Interaction (Phase x Genotype) | F (3, 56) = 1.804 | 0.157 | ns |
| Phase | F (1, 56) = 239.1 | <0.0001 | **** |
| Genotype | F (3, 56) = 2.643 | 0.0581 | # |
| **Dunnett's multiple comparison test** | | | |
| *Dark* | | | |
| double-D2KD vs. AutoD2 | | 0.0208 | * |
| double-D2KD vs. MSN | | 0.9934 | ns |
| double-D2KD vs. Control | | 0.6012 | ns |
| AutoD2 vs. MSN | | 0.0095 | ** |
| AutoD2 vs. Control | | 0.3323 | ns |
| MSN vs. Control | | 0.4363 | ns |
| *Light* | | | |
| double-D2KD vs. AutoD2 | | 0.5879 | ns |
| double-D2KD vs. MSN | | 0.9881 | ns |
| double-D2KD vs. Control | | 0.9093 | ns |
| AutoD2 vs. MSN | | 0.7841 | ns |
| AutoD2 vs. Control | | 0.9329 | ns |
| MSN vs. Control | | 0.9862 | ns |

| Figure | Measurement | # of mice |
|---|---|---|
| S2c | Comparison of controls (Drd2$^{f/w}$, Drd2$^{w/w}$: Dat-cre, A2A-cre, Dat-cre+A2A-cre)<br>Light-Dark Box: Duration in Light Area | n = 5 - 19 |
| S2e-f | Elevated Zero Maze: % Time in open | n = 13 - 38 |
| S3g | Elevated Zero Maze: Number of entries into open | n = 13 - 38 |

| Statistical test | Results | p value | Significance? |
|---|---|---|---|
| **One-way ANOVA** | F (3, 35) = 0.7697 | 0.5188 | ns |
| **Dunnett's multiple comparisons test** | | | |
| control (f/w) vs. control (w/w) (DAT-Cre +/-) | | 0.5345 | ns |
| control (f/w) vs. control (w/w) (A2a-Cre +/-) | | 0.9382 | ns |
| control (f/w) vs. control (w/w) (DAT-Cre +/- ; A2a-Cre +/-) | | 0.9543 | ns |
| **2W ANOVA - Genotype x Age** | | | |
| Interaction (Genotype x Age) | F (3, 173) = 1.540 | 0.206 | ns |
| Genotype | F (3, 173) = 2.982 | 0.0328 | * |
| Age | F (1, 173) = 83.17 | <0.0001 | **** |
| **Fishers Least Significant Difference tes** | | | |
| _12 weeks_ | | | |
| Control vs. double-D2KD | | 0.2937 | ns |
| Control vs. Auto-D2KD | | 0.0685 | # |
| Control vs. MSN-D2KD | | 0.6819 | ns |
| _24 weeks_ | | | |
| Control vs. double-D2KD | | 0.0138 | * |
| Control vs. Auto-D2KD | | 0.0954 | ns |
| Control vs. MSN-D2KD | | 0.0532 | # |
| **2W ANOVA - Genotype x Age** | | | |
| Interaction (Genotype x Age) | F (3, 173) = 0.6243 | 0.6002 | ns |
| Genotype | F (3, 173) = 10.14 | <0.0001 | **** |
| Age | F (1, 173) = 49.37 | <0.0001 | **** |
| **Fishers Least Significant Difference tes** | | | |
| _12 weeks_ | | | |
| Control vs. double-D2KD | | 0.018 | # |
| Control vs. Auto-D2KD | | 0.8228 | ns |
| Control vs. MSN-D2KD | | 0.0016 | ** |
| _24 weeks_ | | | |
| Control vs. double-D2KD | | 0.0016 | ** |
| Control vs. Auto-D2KD | | 0.2962 | ns |
| Control vs. MSN-D2KD | | 0.0014 | ** |

| Figure | Measurement | # of mice | Statistical test |
|---|---|---|---|
| S3c | 15 mg/kg Conditioned Place Preference: Pretest Time on Grid | n = 9 - 10, per contitioned floor, per genotype | **2W ANOVA - Conditioning Floor x Genotype**<br>Interaction (Conditioning Floor x Genotype)<br>Conditioning Floor<br>Genotype |
| S3c | 15 mg/kg Conditioned Place Preference: % Time on Drug-Paired Floor | n = 14 - 20 | **2W mixed-effects model - Genotype x Test**<br>Interaction (Test x Genotype)<br>Test<br>Genotype<br>**Dunnett's multiple comparisons test**<br>*Auto-D2KD*<br>Pretest vs. Test 1<br>Pretest vs. Test 2<br>*MSN-D2KD*<br>Pretest vs. Test 1<br>Pretest vs. Test 2<br>*Control*<br>Pretest vs. Test 1<br>Pretest vs. Test 2<br>*Double-D2KD*<br>Pretest vs. Test 1<br>Pretest vs. Test 2 |

| Results | p value | Significance |
|---|---|---|
| F (3, 67) = 0.1419 | 0.9345 | ns |
| F (1, 67) = 0.006484 | 0.9361 | ns |
| F (3, 67) = 0.1231 | 0.9461 | ns |
| F (5.785, 123.4) = 0.8684 | 0.5171 | ns |
| F (1.928, 123.4) = 45.14 | <0.0001 | **** |
| F (3, 71) = 0.6003 | 0.6169 | ns |
| | 0.0052 | ** |
| | 0.0456 | * |
| | <0.0001 | **** |
| | 0.0014 | ** |
| | 0.0101 | * |
| | 0.0016 | ** |
| | 0.0013 | ** |
| | 0.0061 | ** |

| Figure | Measurement | # of mice | Statistical [illegible] | Results | p value | Significance [illegible] |
|---|---|---|---|---|---|---|

| Figure | Measurement | # of mice | Statistical [illegible] | Results | p value | Significance [illegible] |
|---|---|---|---|---|---|---|
